# The elemental mechanism of transcriptional pausing

**DOI:** 10.1101/422220

**Authors:** Jason Saba, Xien Chua, Tatiana V. Mishanina, Dhananjaya Nayak, Tricia A. Windgassen, Rachel A. Mooney, Robert Landick

## Abstract

Transcriptional pausing underpins regulation of cellular RNA biogenesis. A consensus pause sequence that acts on RNA polymerases (RNAPs) from bacteria to mammals halts RNAP in an elemental paused state from which longer-lived pauses can arise. Although the structural foundations of pauses prolonged by backtracking or nascent RNA hairpins are recognized, the fundamental mechanism of the elemental pause is less well-defined. Here we report a mechanistic dissection that establishes the elemental pause signal (*i*) is multipartite; (*ii*) causes a modest conformational shift that puts RNAP in an off-pathway state in which template base loading but not RNA translocation is inhibited; and (*iii*) can easily enter pretranslocated and one-base-pair backtracked states despite principally occupying the half-translocated state observed in cryo-EM structures of paused RNAPs. Our findings provide a mechanistic basis for the elemental pause and a framework to understand how pausing is modulated by sequence, cellular conditions, and regulators.

## INTRODUCTION

During the first step in gene expression, transcription by RNA polymerase (RNAP) at ∼30 nt/s or faster is interrupted by ≥1-s pause events every 100-200 bp (Landick, 2006; Larson *et al.*, 2014; Chen *et al.*, 2015). These pauses underpin diverse mechanisms that regulate gene expression in both prokaryotes and eukaryotes (Figure 1A), including attenuation, antitermination, and promoter-proximal pausing (Jonkers and Lis, 2015; Zhang and Landick, 2016; Mayer *et al.*, 2017). Pausing also couples transcription to translation in bacteria or to mRNA splicing in eukaryotes (Landick *et al.*, 1985; Proshkin *et al.*, 2010; Mayer *et al.*, 2017); defines temporal and positional windows for binding of small molecules, regulatory proteins, or regulatory RNAs to the nascent RNA transcript (Wickiser *et al.*, 2005; Artsimovitch and Landick, 2002); mediates nascent RNA folding (Pan *et al.*, 1999; Pan and Sosnick, 2006; Steinert *et al.*, 2017); and enables termination (Gusarov and Nudler, 1999; Proudfoot, 2016). Conversely, RNA folding and the interactions of cellular molecules and complexes (*e.g*., ribosomes) with the elongating transcription complex (EC) modulate pausing (Toulokhonov *et al.*, 2001; Artsimovitch and Landick, 2002; Yakhnin *et al.*, 2016; Zhang and Landick, 2016). Despite its crucial role in cellular information processing, the biophysical mechanism of pausing remains incompletely defined.

**FIGURE 1.**
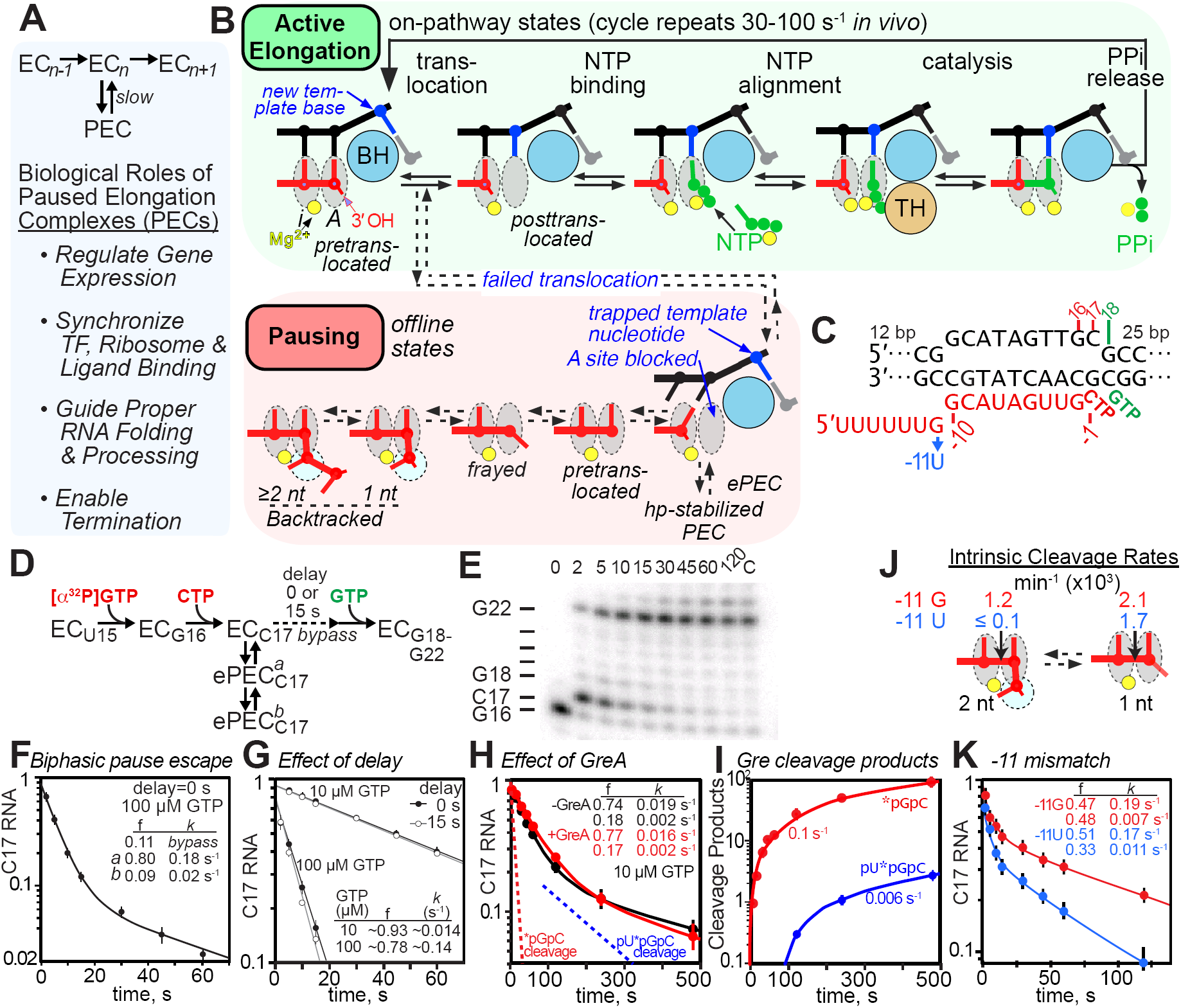
The consensus sequence elemental pause. (**A**) The simplest elemental pause kinetic scheme and the biological roles of pausing. (**B**) Model of RNAP active site during the normal nucleotide addition cycle (*green*) consisting of RNA-DNA translocation, NTP substrate binding, active-site closing and NTP alignment (mediated by the TL and BH), phosphoryl transfer (catalysis), and PPi release. Catalysis is reversible when PPi remains present. Pausing (*red*) occurs when altered RNAP-DNA-RNA interactions create a barrier to completion of translocation that blocks entry of the next template nucleotide into the active site, creating the elemental pause state. The elemental pause state can rearrange further into backtracked or hairpin-stabilized pauses depending on other RNA-DNA sequences. (**C**) Central region of the RNA:DNA scaffold used for pause assays (complete scaffold is in Figure S1A). The -11U mismatch substitution was used to test possible contributions of backtracking (see panels **K** and **J**). (**D**) Pause assay scheme consistent with biphasic pause escape intermediates. (**E**) Example of pause assay products separated by denaturing gel electrophoresis (radiolabeled by ^32^P-G16 incorporation). (**F**) Observed levels of C17 RNA as a function of time in a simple elemental pause assay at 37 °C and 100 µM GTP. (**G**) Effect of delayed addition of 10 µM or 100 µM GTP after formation of C17 PECs. (**H**) Effect of GreA and GreB (1 µM each) on elemental pausing at 10 µM GTP. (**I**) Rate of appearance of GreA- and GreB-induced cleavage products during pause assay in panel H (see Figure S3A,B,C,D for measurements of GreA/B cleavage rates and effects on pausing). (**J**) Rates of intrinsic cleavage if C17 halted ePECs matched with pause assays shown in panel K. The -11U substitution reduced intrinsic cleavage in the -1 backtrack register be a factor of >50 (see Figure S3E,F for measurement of intrinsic cleavage rates). (**K**) Effect of - 11 U mismatch on the elemental pause at 100 µM GTP with pause prone-RNAP.

Multiple pause mechanisms exist, but most pauses that mediate gene regulation are triggered initially by sequence-specific interactions of DNA and RNA with RNAP. An increasingly accepted view is that these initial interactions interrupt the nucleotide addition cycle by promoting entry of RNAP into a state termed the elemental pause (Landick, 2006), creating an elemental paused elongation complex (ePEC; Figure 1B). The ePEC can then rearrange into long-lived pause states by backtracking (reverse translocation of RNA and DNA), by pause hairpin (PH) formation in the RNA exit channel that alters RNAP conformation (at least in bacteria), or by interactions of diffusible regulators with the ePEC. Recent cryoEM structures of artificially assembled PECs suggest the ePEC is half-translocated with a tilted RNA-DNA hybrid, meaning that the RNA but not the DNA is translocated and the next template base is still sequestered in the downstream DNA duplex. The hairpin-stabilized PEC is additionally inhibited by a rotation of the swivel module (including the clamp, shelf, and SI3) that inhibits trigger-loop folding (Kang *et al.*, 2018a; Guo *et al.*, 2018).

High-throughput sequencing of nascent RNAs from bacteria (NET-seq) reveals a consensus elemental pause sequence conserved among diverse bacterial RNAPs and mammalian RNAPII whose effects on pausing *in vitro* are consistent with a block in template DNA translocation sometimes accompanied by modest backtracking (Larson *et al.*, 2014; Vvedenskaya *et al.*, 2014; Imashimizu *et al.*, 2015). Although earlier work establishes contributions of multiple pause signal components (upstream RNA, RNA:DNA hybrid, downstream fork-junction, and downstream DNA) to hairpin-stabilized pausing (Chan and Landick, 1993; Wang *et al.*, 1995; Chan *et al.*, 1997), the definition of the consensus elemental pause signal has varied and it is unknown if the discrete components affect a common step in the elemental pause mechanism.

Additionally, questions remain about the structure and properties of the ePEC. A longstanding debate is whether the ePEC is an on-pathway state unable to translocate DNA or RNA in a largely unchanged RNAP due to the thermodynamic properties of the RNA-DNA scaffold (*i.e*., a pretranslocated pause; (Bai *et al.*, 2004; Bochkareva *et al.*, 2012) or if it represents an offline state generated by conformational rearrangement of RNAP that forms in kinetic competition with the on-pathway steps (Landick, 2006; Herbert *et al.*, 2006; Kireeva and Kashlev, 2009; Imashimizu *et al.*, 2013; Kang *et al.*, 2018a). Uncertainty also exists as to whether the elemental pause is non-backtracked (Landick, 2006; Herbert *et al.*, 2006; Kireeva and Kashlev, 2009; Kang *et al.*, 2018a) or must be backtracked one or more registers (Dangkulwanich *et al.*, 2013; Forde *et al.*, 2002; Galburt *et al.*, 2007; Mejia *et al.*, 2014; Tadigotla *et al.*, 2006; Maoileidigh *et al.*, 2011).

To address these questions, we combined kinetic analyses of pausing using elemental pause sequence variants or mutant RNAPs with precise structural probes of translocation, trigger-loop folding, and clamp conformation. Our results lead us to propose a multistate model of elemental pausing in which template-base loading in a half-translocated offline intermediate limits pause escape.

## RESULTS

### The ePEC is an offline state formed in a branched kinetic mechanism

To probe the elemental pause mechanism, we used kinetic analyses of ECs reconstituted on a synthetic RNA-DNA scaffold encoding a consensus elemental pause sequence (Figure 1C and Figure S1A; Larson *et al.*, 2014). We first asked if the pause signal always causes ECs to bifurcate into paused and rapidly elongating (bypass) fractions. We measured C17 pause RNA as a function of time when radiolabeled G16 ECs were extended with 100 µM each CTP and GTP (Figure 1D,E,F). Most ECs (≳80% at 37 °C) entered a paused state (lifetime ≳5 s; *a*, Figure 1D,F), whereas some ECs (≳20%) transcribed past C17 rapidly (lifetime ≳0.1 s; *bypass*, Figure 1D,F). Invariably, a minor ePEC population (typically ≳15%) escaped more slowly (lifetime ≳ 100 s; *b*, Figure 1D,F), requiring double-exponential fitting of the escape rate. Pause bypass was evident from y-intercepts <1 in the double-exponentials fits (Figure 1F). Interestingly, the amounts of slow ePEC and bypass fractions varied among RNAP preparations (Figure S1B). This kinetic malleability of ePECs differed from the reproducible kinetics typically observed for hairpin-stabilized pauses like the *his*PEC (Toulokhonov *et al.*, 2001; Kyzer *et al.*, 2007; Kang *et al.*, 2018a).

To confirm that the ePEC is an offline state formed in competition with bypass nucleotide addition (*i.e*., in a branched kinetic mechanism; Figure 1D), we performed three additional tests. First, we asked whether the fraction of ECs entering the ePEC state increased if GTP (the NTP required for pause escape) were withheld to allow more time for ePEC formation. Unlike for the hairpin-stabilized *his*PEC (Toulokhonov *et al.*, 2007), halting ECs at C17 for 15 s before GTP addition did not change the 10-20% EC that bypassed the pause (Figure 1G). However, the bypass fraction was shifted from ∼22% at 10 µM GTP to ∼7% at 100 µM GTP, suggesting that GTP can affect a partition between the on-pathway EC and offline elemental pause state. Second, using data from Larson *et al*., 2014, we found that the bypass fraction varied but appeared to plateau at high GTP (Figure S1C). Because extrapolation of bypass from bi-exponential fits is an approximation, we next used numerical integration of rate equations as a third test to ask if reaction progress curves are better fit by an unbranched or branched kinetic mechanism at saturating GTP (10 mM; Figure S1A,F; Methods). The unbranched (on-pathway pause) mechanism did not fit the data well, thus favoring a branched mechanism (Figure S1F).

We also used kinetic modeling to reexamine the proposed on-pathway pausing reported by Bochkareva *et al*. (2012). Using their optimal pause scaffold sequence, we replicated the reported high-efficiency pausing (Figure S1D,E). However, this scaffold positions the downstream DNA end (+17) within RNAP, so that forward translocation reduces downstream DNA-RNAP interaction. Consistent with a prior finding that downstream DNA truncation increases pausing (Kyzer *et al.*, 2007), pause bypass was readily detected when the downstream DNA end was positioned outside RNAP as in natural ECs (Figure S1D,E). This result highlights why highly efficient pauses, even if they isomerize to an offline state, may appear to be on-pathway when bypass falls below ∼5%. We confirmed the branched kinetic pause mechanism on the Bochkareva *et al*. scaffold at saturating GTP by kinetic modeling (Figure S1G). In agreement with most prior ensemble (Kassavetis and Chamberlin, 1981; Kyzer *et al.*, 2007; Toulokhonov *et al.*, 2007; Strobel and Roberts, 2015) and single-molecule (Herbert *et al.*, 2006; Larson *et al.*, 2014; Gabizon *et al.*, 2018) analyses of pausing, we conclude that the elemental pause is an offline state that forms in competition with on-pathway nucleotide addition (*i.e*., in a branched kinetic mechanism).

### The consensus ePEC readily backtracks by one bp but reversal of one-bp backtracking does not limit the rate of pause escape

The existence of a minor, variable, slowly escaping ePEC population has been a consistent and puzzling feature of pause assays using synthetic scaffolds (Figure 1F). We wondered if the slow pause population could be explained by backtracking, an EC conformational change, or a subpopulation of chemically altered RNAP. Variation of the slow ePEC fraction among RNAP preparations (Figure S1B) seemed to favor a chemically altered subpopulation. We found that absence of neither the ω subunit nor the αCTDs changed the slow ePEC fraction (not shown). More definitively, transcription through tandem consensus pause sequences revealed formation of the slow fraction from all RNAPs arriving at the second pause site rather than a chemically altered, slow subpopulation of RNAP that would be filtered out by the first pause site (Figure S2).

We next used GreA- or GreB-induced cleavage to ask if the minor slow ePEC fraction was backtracked. Either or both GreA or GreB had little effect on pause lifetimes, even though 2-nt cleavage products, indicative of a 1-bp backtracked state, appeared much faster than the rate of ePEC escape and 3-nt cleavage products, indicative of a 2-bp backtracked state, were detectable (Figure 1H,I; Figure S3B&D). Thus, even though the scaffold structure disfavors ≥2 bp backtracking due to a -12 rU-dG mismatch, ePECs may shift from the 1-bp backtracked state to ≥2 bp backtracked states, at least in the presence of GreA/B. We next tested the effect on both pausing and intrinsic cleavage of a -11rU-dC mismatch that would disfavor even 1-bp backtracking using an RNAP that formed a high level of slow fraction (Figure 1C,K,J; Figure S3E,F). The -11rU-dC mismatch virtually eliminated ≥2-nt cleavage indicative of backtracking, reduced 1-nt cleavage indicative of the pretranslocated register, and modestly decreased the slow pause fraction.

The relative ease with which ePECs entered backtracked registers is notable given the half-translocated state observed in an ePEC cryo-EM structure (Kang *et al.*, 2018a; Guo *et al.*, 2018). Backtracking of ePECs is detectable both *in vivo* and *in vitro* (Larson *et al.*, 2014; Imashimizu *et al.*, 2015), but the half-translocated or pretranslocated states appears to be predominant (Figure 1J; Larson *et al.*, 2014). Our GreA/B and intrinsic cleavage results establish that slow escape from a 1-bp backtrack state explains neither the major nor the minor pause fractions even though the 1-bp backtrack state is accessible. A parsimonious explanation is that the ePEC readily equilibrates among several active-site states, including half-translocated, pretranslocated, and backtracked, with the half-translocated state being the dominant species in which the kinetic block to pause escape is manifest. The slow pause state may be backtracked by ≥2-bp (since 0.006 s^-1^ 3-nt cleavage is close to the 0.002s^-1^ escape rate; Figure 1H,I), but its persistence in the -11U mutant suggests other changes to the ePEC must also contribute (see Discussion).

### The elemental pause signal is multipartite

Available NET-seq data have been interpreted to suggest an elemental pause signal involving just the upstream fork-junction (usFJ and downstream fork-junction (dsFJ) sequences or a multipartite signal that additionally depends on the hybrid (Hyb) and downstream DNA (dsDNA) sequences (Larson *et al.*, 2014; Vvedenskaya *et al.*, 2014; Imashimizu *et al.*, 2015). To test the proposed roles of Hyb and dsDNA sequences and additionally to ask if the different pause signal elements combine additively to define a single energetic barrier to pause escape, as observed for the hairpin-stabilized *his* pause signal (Wang *et al.*, 1995; Chan *et al.*, 1997), we analyzed effects on pausing of substitutions in each element separately and in combination (Figure 2A,B).

**FIGURE 2.**
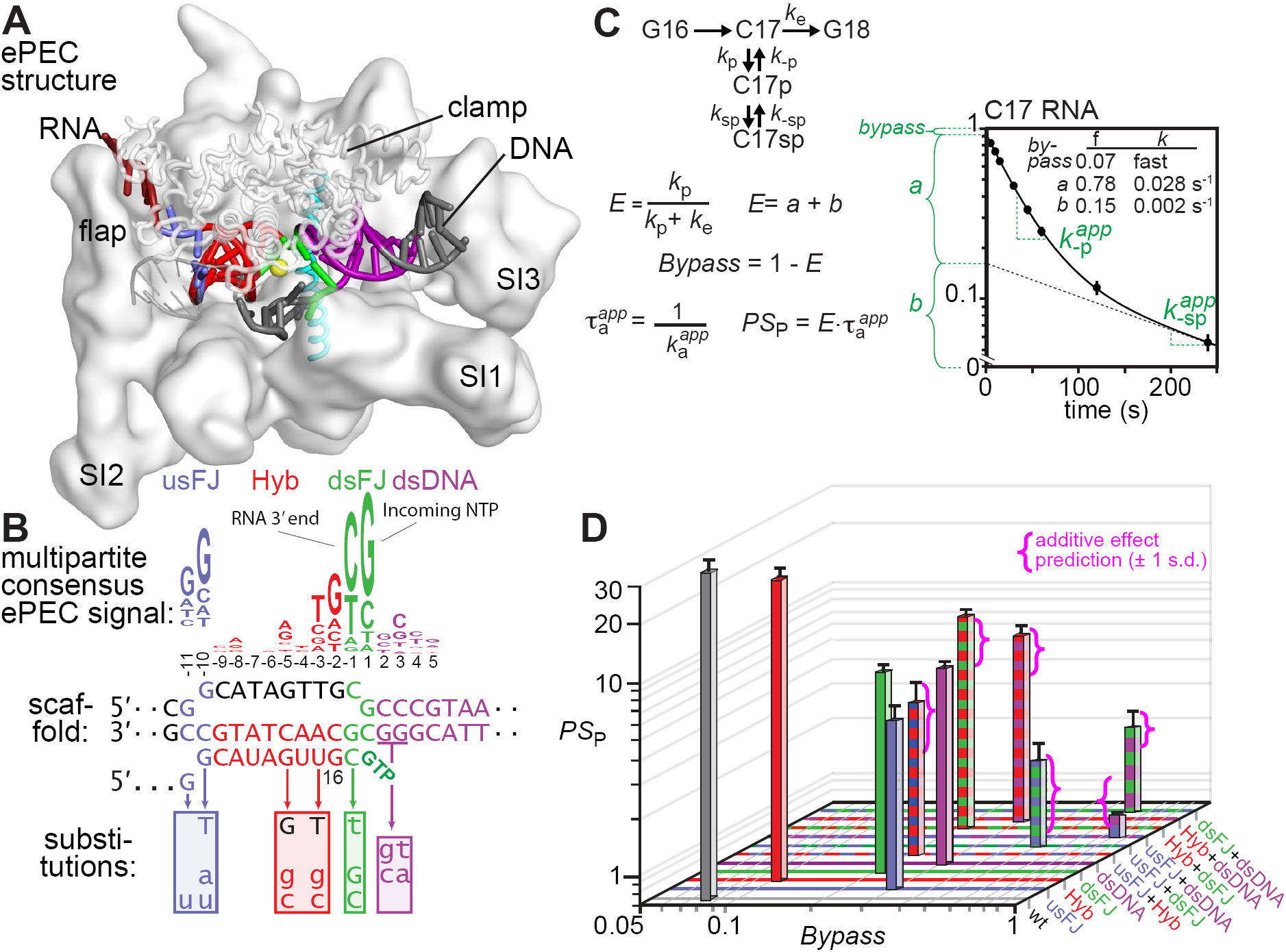
The elemental pause signal is multipartite. (**A,B**) The consensus ePEC. The ePEC structure (pdb 6bjs; Kang *et al.*, 2018a) is shown above the consensus pause sequence (Larson *et al.*, 2014) color-coded as usFJ (blue), Hyb (red), dsFJ (green), and dsDNA (purple). Scaffold substitutions tested for effects on pausing are shown below the location of the changes. (**C**) Two component pause mechanism, equations used to calculate pause efficiencies and pause strengths, and example plot. The pause assay shown is for the “wild-type” (unsubstituted) consensus pause scaffold at 37 °C and 10 µM GTP. Pause fractions (fast fraction *a*, slow fraction *b*) and escape rates k_-p,app_ and k_-sp,app_ were obtained by nonlinear regression using a double-exponential equation. (**D**) Plot of pause strength (PS_P_) *vs.* bypass for each scaffold variant. Combination are indicated by “+” (*e.g*., usFJ + Hyb). The magenta brackets show the predicted range of *PS*_P_ values (95% confidence interval) for combinations of substitutions in individual pause signal elements assuming the pause signal elements additively and independently affect a single energy barrier to pause escape (additive effects on ΔG^‡^ are multiplicative in *PS*_P_). Note that a single RNAP preparation was used for all experiments shown in this figure.

We measured pause lifetimes and the bypass, pause, and slow pause fractions using two-exponential fits of C17 RNA *vs.* time at 10 µM GTP (Figure 2C). Because a translocation barrier would affect both pause formation and escape, we plotted pause strength (PS; pause efficiency times the lifetime [τ] of the major pause species) vs. pause bypass (Figure 2D). Alone, the usFJ, dsFJ, and dsDNA mutants reduced pause strength by a factor of ∼5, whereas our Hyb mutant had an ∼1.4-fold effect (but see larger effects in (Bochkareva *et al.*, 2012). Combinations of mutants gave additive effects on pause strength (Figure 2D, magenta brackets; additive effects on an energy barrier are multiplicative in τ or PS; *e.g.*, reduction of PS by factors of 1.4 and 3.7 for the Hyb and dsDNA substitutions predicts a combined reduction by a factor of 5.1 *vs*. the factor of 5.2 observed). Additive effects are expected if pause signal components independently affect the same energetic barrier to pause escape (*e.g*., translocation of the template base).

The lifetimes of the major and minor pause states were highly correlated for the pause-sequence variants (Figure S4B). In contrast, the slow pause/pause ratio varied from 0.04 (usFJ) to 0.32 (Hyb-dsDNA) and was uncorrelated with either the total ePEC fraction or the lifetime of the major pause species (Figure S4C). These results are consistent with a model in which both the major pause species and the minor, slowly escaping pause species pass through a common barrier for pause escape (*e.g.*, template-base loading). The significant variation in the amount of slow pause species provides additional evidence that the slow species arises by kinetic partitioning of a single RNAP population and not from a chemically distinct “slow” subpopulation of RNAP.

We conclude that the elemental pause signal is indeed multipartite with significant contributions from sequences in the usFJ, Hyb, dsFJ, and dsDNA to a common kinetic barrier to escape from multiple paused states.

### Hybrid translocation is not rate-limiting for elemental pause escape

To ask if the barrier to elemental pause escape corresponds to the half-translocated intermediate identified by cryoEM (Kang *et al.*, 2018a; Guo *et al.*, 2018), we next tested for ePEC translocation at the usFJ and dsFJ using a fluorescence-based translocation assay developed by Belogurov and co-workers (Figures 3A and 4A; Malinen *et al.*, 2012). In this assay, the fluorescent guanine analog 6-methylisoxanthopterin (6-MI) is positioned in the template DNA (usFJ) or nontemplate DNA (dsFJ) so that an adjacent guanine unstacks from 6-MI upon translocation of the hybrid or the dsDNA; this unstacking increases 6-MI fluorescence (Figure 3A). We compared translocation of the hybrid or the dsDNA leading to increased 6-MI fluorescence on ePEC scaffolds to a control scaffold previously found to translocate rapidly upon 3’-CMP addition (Figure 3B; Malinen *et al.*, 2012; Hein *et al.*, 2014). Because both the control and ePEC scaffold encode C as the RNA 3’ nucleotide and G as the next nucleotide after translocation (Figures 3B,C and 4A,B), differences in their behavior can be attributed to the usFJ, Hyb, and dsDNA. To calibrate the 6-MI signal change from the posttranslocated hybrid, we compared the effects of CMP, 3’dCMP, or 2’dCMP (Figure 3D,E). A 3’deoxy RNA shifts the hybrid toward the pretranslocated register, whereas 2’deoxy RNA shifts the hybrid toward the posttranslocated register (Malinen *et al.*, 2014). For both scaffolds, 2’dCMP gave a 6-MI signal increase comparable to CMP, whereas 3’dCMP reduced the signal increase. These results suggested that addition of CMP shifts both hybrids toward the posttranslocated register, although the absolute fluorescence change for the ePEC scaffold was about half that of the control scaffold. Rapid-quench and stopped-flow kinetic measurements revealed that CMP added rapidly (∼300 s^-1^) followed by a rapid (∼200 s^-1^) rise in fluorescence signal indicating translocation on both scaffolds. In contrast, subsequent extension to G18 of the ePEC but not the control EC was slow (∼0.2 s^-1^ *vs*. 40 s^-1^; Figures 3D and 3E).

**FIGURE 3.**
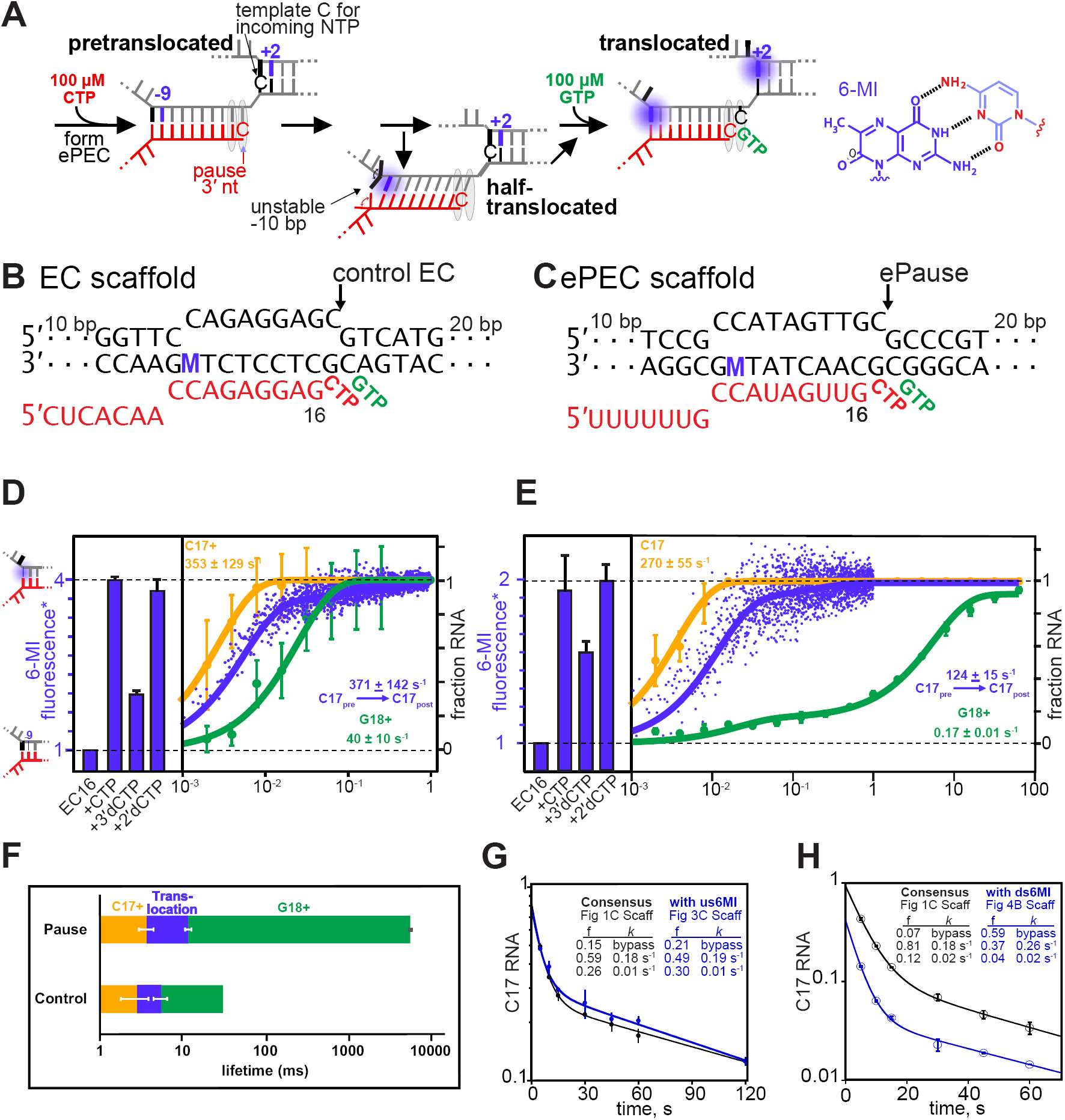
Translocation of the RNA:DNA hybrid is not rate-limiting for elemental pause escape. (**A**) Scheme for translocation following CTP addition to control EC or ePEC scaffolds. The locations of 6-MI for both usFJ and dsFJ probes are indicated, but a probe was present in only one location in the scaffolds used (see panels B and C and Figures 4A and 4B). 6-MI is a GMP base analog that fluoresces unless quenched by stacking with an adjacent GMP. (**B** and **C**) Scaffolds used to reconstitute control EC and ePEC for fluorescence experiments. M, position of 6-MI. (**D**) Equilibrium fluorescence changes (blue bars) upon addition of incoming NTP or NTP analog to the non-pausing EC scaffold measured at 37 °C. Nucleotide addition (orange and green traces) and translocation (blue trace) rates were determined using KinTek quenched-flow (RQF-3) or stopped-flow (SF-300X) instruments, respectively. C17+ represents the fraction of RNA at or beyond C17. G18+ represents the fraction of RNA at or beyond G18. 6-MI fluorescence normalized to that observed after 2’dCMP addition (post-translocation bias). (**E**) Experiments performed on an ePEC scaffold as in D. (**F**) Data from **D** and **E**. Mean times for CMP addition, translocation, and subsequent GMP for the ePEC and control EC. (**G**) Pause kinetics at 100 µM GTP for the 6-MI-containing ePEC scaffold compared to the consensus pause scaffold. f, kinetic fractions. *k*, rate constants for pause fractions.

Because the 6-MI and -10 substitutions in the ePEC scaffold could affect pausing, we verified that the 6-MI ePEC scaffold gave pause fractions (∼85%) and lifetimes similar to the wild-type consensus scaffold (Figure 3G,H). Thus, the ePEC fluorescence signal could not be explained by the ∼15% bypass fraction and must have arisen in significant part from the paused species. We also verified that CMP addition occurred fully in the fluorescence assay (Figure S5). We conclude that the ePEC hybrid translocates rapidly after CMP addition, resembling the control scaffold. The reduced level of unquenching (∼50% of the control scaffold) could reflect lower 6-MI fluorescence in a half-translocated hybrid, since the ePEC cryoEM structure suggests -10 dC (*vs*. -10 dG in the assay scaffold) remains at least partially paired in the hybrid with an altered interaction with the lid (Kang *et al.*, 2018a). Alternatively, -10 dG may unstack in our assay at 37 °C but the pretranslocated and half-translocated hybrids may be in dynamic equilibrium (consistent with intrinsic hydrolysis readout; Figure 1J). These results also confirm that most ePECs are not backtracked, which would not unquench 6-MI (Figures 1I and S3). Together the data are fully consistent with a view that the ePEC, once formed, may easily fluctuate among multiple translocation states.

### Downstream DNA translocation is rate-limiting for elemental pause escape

To test whether dsDNA fails to translocate in the ePEC (as predicted from the ePEC structure), we used an ePEC scaffold for which translocation of the +1 dC-dG bp would generate a +2 6-MI fluorescence signal (+1 dG would shift into an RNAP pocket that aids pause escape; (Vvedenskaya *et al.*, 2014). Placement of a dT-dA bp at +3 was necessary to generate a strong +2 6-MI signal. The changes needed for the 6-MI assay weakened the elemental pause signal but still allowed ECs to enter the pause state (Figure S3H), possibly to greater extent prior to GTP addition (*e.g*., see effect of GTP in Figure 1G). For the control scaffold, addition of CMP or 2’dCMP or extension with CTP+GTP to G19 gave a large 6-MI signal indicating translocation after C17 nucleotide addition. In contrast, the ePEC scaffold gave minimal 6-MI signal after CMP or 2’dCMP addition (consistent with translocation in only a bypass fraction) compared to extension to G19, which produced a strong 6-MI signal. These data confirm that the dsDNA in the ePEC does not translocate. Since the dsDNA in the ePEC did not translocate, we used GTP binding to 3’dCMP-ePEC and control 3’dCMP-EC to assess translocation potential. For the control EC, a clear increase in 6-MI signal was evident as GTP bound in the active site with apparent *K*_d_ ≈ 400 µM (Figure 4D, inset). This signal, which was reduced by interference from the high levels of GTP present, indicated that the post-translocated register was stabilized by GTP binding to the control EC (Figure 4C; (Malinen *et al.*, 2014). However, GTP binding was unable to shift the 3’dC17 ePEC to the post-translocated register (Figure 4D). We conclude that even the weakened elemental pause signal in the dsFJ ePEC assay scaffold prevented dsDNA translocation and thus template-base loading. Taken together, the usFJ and dsFJ 6-MI data confirm that the half-translocated state detected in ePEC cryoEM structure also forms in actively transcribing complexes, and that template-base loading is the relevant barrier to nucleotide addition in the ePEC.

**FIGURE 4.**
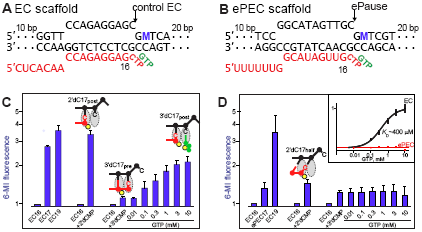
Translocation of the incoming template DNA limits elemental pause escape. (**A** and **B**) Scaffolds used to reconstitute control EC and ePEC for fluorescence experiments. M, position of 6-MI. (**C** and **D**) Equilibrium fluorescence changes of control EC and ePEC, respectively, upon reaction with NTP or NTP analog or binding of GTP to 3’deoxyC-containing complexes. RNA extension was verified by quantifying radiolabeled RNAs (see Figure S5). Inset, GTP-binding curve fit for the 3’dC17 EC. The 3’dC-ePEC has inhibited GTP binding (flat, dotted red line). Error bars are SD of triplicate measurements.

### NTP binding rather than TL folding limits escape from the elemental pause

As a second approach to interrogate the status of the RNAP active site in the ePEC, we tested the effect of the consensus pause sequence on formation of disulfide bonds between Cys substitutions engineered to report TL conformation (Cys-pair reporters, CPRs; Nayak *et al.*, 2013). Previous studies established that a β’ 937 Cys substitution in the trigger helix near the active site crosslinks efficiently to a Cys substitution at β’ 736 when the trigger helices formed (folded CPR, F937-736; Figure 5A). Other CPRs (P937-687, U937-1135, or U937-1139) report the partially folded or unfolded conformations of the TL (Figure 5A). The CPRs are sensitive to TL conformation when oxidized by cystamine (CSSC) because the CPR disulfide competes with formation of mixed disulfides (Figure 5B). These CPRs combined with other probes and a cryoEM structure revealed that the hairpin-stabilized *his*PEC fails to add the next nucleotide because it forms a swiveled PEC conformation that inhibits TL folding (Hein *et al.*, 2014; Nayak *et al.*, 2013; Kang *et al.*, 2018a). In contrast, the ePEC, whose cryoEM structure is not swiveled, did not exhibit a restricted TL conformation (compare to control EC; Figure 4C,D,E). If anything, the ePEC accessed the folded TH conformation more readily than did the control EC, consistent with ePEC access to the pretranslocated register (Figure 1J) that is thought to increase TL folding (Malinen *et al.*, 2014; Liu *et al.*, 2016). Thus, TL folding does not appear to limit escape from the elemental pause.

**FIGURE 5.**
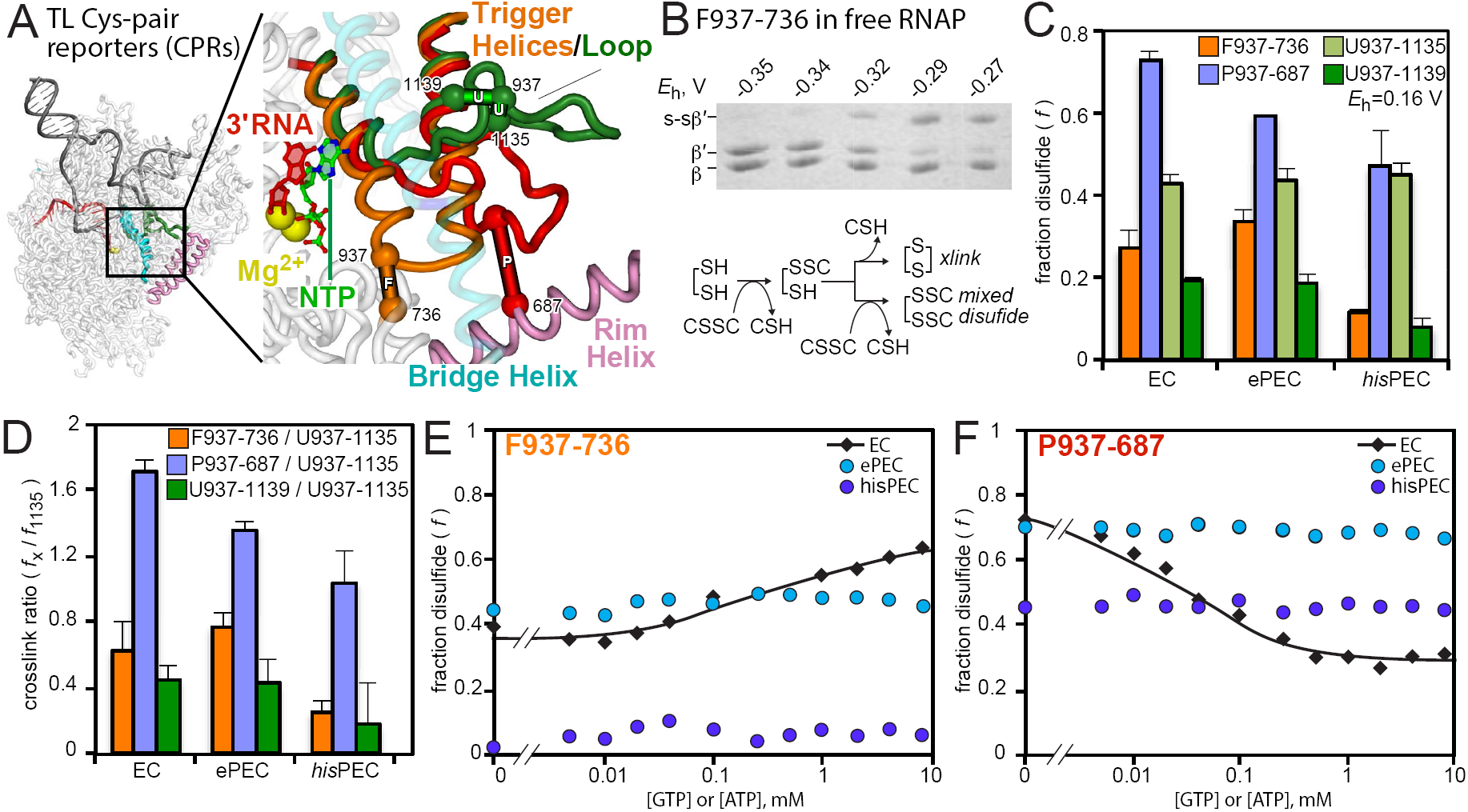
NTP binding but not TL folding is inhibited in the ePEC. Scaffolds used for these experiments are shown in Figure S6. (**A**) Location of F937-736, P937-687, U937-1137, and U937-1139 Cys-pair reporters to test various conformations of the TL. The SI3 domain inserted in the *Eco*RNAP TL is not shown. A complete description of these reporters is published elsewhere (Nayak *et al.*, 2013; Windgassen *et al.*, 2014). (**B**) Example CPR reaction showing SDS-PAGE mobility of uncrosslinked and crosslinked β’ as a function of redox potential generated by increasing concentrations of cystamine. Reaction scheme for crosslink generation by cystamine. (**C**) Fraction crosslink formed by CPRs for the EC, ePEC, and hairpin-stabilized *his*PEC in which TL mobility is restricted. (**D**) Ratio of crosslinks to the U1135 crosslink. A high ratio indicates a greater (**E**) F937-736 crosslink as a function of GTP or ATP concentration in 3’dC or 3’dU containing EC, ePEC, and *his*PEC. (**F**) P937-687 crosslink as a function of GTP or ATP concentration in 3’dC or 3’dU containing EC, ePEC, and *his*PEC.

Incorporation of a 3’-dNMP in ECs and PECs allows the CPRs to detect TL-folding stimulated by NTP binding in the EC and its inhibition in the *his*PEC (Nayak *et al.*, 2013). Consistent with results of the 6-MI dsFJ translocation assay, F937-736 and P937-687 crosslinking in 3’dCMP-ePEC, and therefore TL conformation, were unaffected by high GTP concentration even though the crosslinks formed readily. In contrast, CPR crosslinking in control 3’dCMP-EC exhibited an obvious shift toward TL folding upon ATP binding (Figure 5E,F; Nayak *et al.*, 2013). These results are consistent with the ePEC cryoEM structure and translocation assay results, supporting a view that inability to load the template base inhibits NTP binding in the ePEC.

### Clamp loosening but not extensive clamp opening accompanies elemental pausing

The role of clamp conformation in elemental pausing is uncertain. Although a crystal structure of *Tth*RNAP on a partial ePEC scaffold suggested the clamp could open in the ePEC (Weixlbaumer *et al.*, 2013), more recent cryoEM structures and biochemical probing suggested the clamp remains closed in the ePEC but can swivel upon pause hairpin formation (Guo *et al.*, 2018; Kang *et al.*, 2018a). To probe the role of clamp conformation in elemental pausing, we examined the effect of stabilizing the clamp in the closed (unswiveled) conformation using an engineered disulfide bond between the lid and flap (β’258i-β1044; Figures 6A and 6B; Kang *et al.*, 2018a). In contrast to suppression of pausing for the hairpin-stabilized *his*PEC (Figure 6C; Hein *et al.*, 2014; Kang *et al.*, 2018a), the closed-clamp disulfide had minimal effect on the ePEC lifetime (Figure 5D). We conclude that, unlike for hairpin-stabilized pausing and consistent with the cryoEM structure, clamp swiveling (or opening) is not required in the ePEC.

**FIGURE 6.**
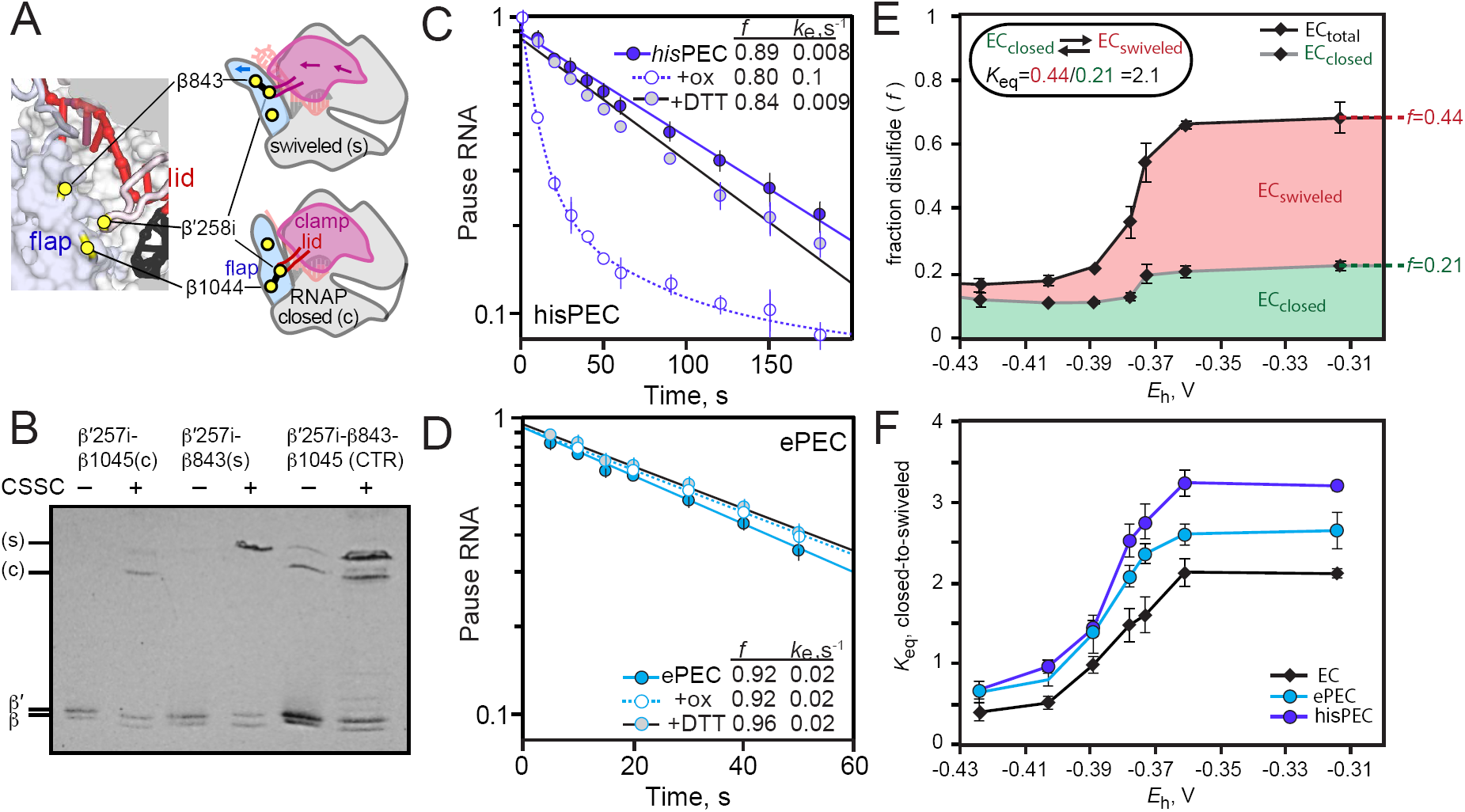
Restriction of clamp movement has less effect on ePEC than on hairpin-stabilized PEC. (**A**) Location of disulfides used to restrict clamp movement or generate the Cys-triplet reporter (CTR; described in Hein *et al.*, 2014; Kang *et al.*, 2018b). (**B**) Example β-β’ disulfide mobilities during SDS-PAGE illustrating the identification of the closed (β1044-β’258i) or swiveled (β843-β’258i) disulfides. (**C**) Effect of the closed clamp crosslink on pausing by the hairpin-stabilized *his*PEC (Scaffold shown in Figure S6E). (**D**) Effect of the closed clamp crosslink on pausing by the ePEC (scaffold shown in Figure S1A). (**E**) CTR assay on an EC scaffold illustrating the formation of the closed and swiveled crosslinks as a function of redox potential generated by increasing concentrations of cystamine (scaffold shown in Figure S6F). (**F**) CTR assay of clamp swiveling in EC, ePEC, and *his*PEC (scaffolds shown in Figure S6F, Figure S6G, and Figure S6H, respectively).

Even though full clamp swiveling is not required for elemental pausing, we wondered if any change in clamp conformation accompanied formation of the ePEC. To investigate this question, we used a variant of the disulfide bond probing strategy in which three Cys residues were positioned in RNAP such that either the closed-clamp disulfide (β’258i-β1044) or the swiveled-clamp disulfide (β’258i-β843) could form (Kang *et al.*, 2018a). This Cys triplet reporter (CTR) enabled a convenient measure of the energetic balance between closed and swiveled clamp conformations because the different crosslinked β’-β polypeptides could be readily distinguished by denaturing gel electrophoresis (Figure 6B; Kang *et al.*, 2018a). The ratio of closed-to-swiveled crosslinks shifts during oxidation with cystamine likely because the mixed disulfide intermediates destabilize the closed-clamp conformation (Figure 6E). Thus, cystamine oxidation shifts the unswiveled-to-swiveled equilibrium toward the swiveled conformation, making it a sensitive assay of clamp energetics. This shift is greater for the *his*PEC (which favors clamp swiveling; Kang *et al.*, 2018a), but intermediate between the EC and *hisPEC* for the ePEC (Figure 6F). We conclude that formation of the ePEC is accompanied by a loosening of clamp contacts.

### A conserved Arg in Fork Loop 2 may help inhibit DNA translocation at an elemental pause

We wondered if specific amino acids in RNAP inhibit template base translocation in the ePEC. Two candidates were β’K334 and βR542 (Figure 7A). β’K334 in switch 2 contacts the template DNA backbone adjacent to a dC blocked from active-site loading in half-translocated *Sce*RNAPII EC and *Tth*EC crystal structures (Brueckner and Cramer, 2008; Weixlbaumer *et al.*, 2013) and the ePEC cryoEM structure. βArg542 in fork loop 2 appears to contact the template +1dC in the ePEC cryoEM structure. Although the modest resolution of the cryoEM structure makes this assignment tentative, interaction of this conserved Arg with a pretranslocated template +1C also is seen in the half-translocated *Tth*RNAP crystal structure (Weixlbaumer *et al.*, 2013), in a crystal structure of a *Tth*RNAP open complex formed on the *B. subtilis pyrG* promoter (Murakami *et al.*, 2017), and in a 1-bp backtracked *Tth*RNAP PEC (Sekine *et al.*, 2015). To ask if either β’K334 or β542 played a key role in inhibiting template-base loading in the ePEC, we generated Ala substitution mutants and compared their pausing behaviors to wild-type RNAP. Because the strong consensus pause sequence could mask the effect of a single amino-acid contact, we also assayed the Ala mutants on the usFJ and dsFJ mutant templates (Figure 2B). Strikingly, β’K334A resembled the wild-type enzyme on both consensus and mutant pause scaffolds, whereas βR542A decreased pausing by a factor of 2 on the consensus pause and dsFJ templates, but significantly more on the usFJ template (13X effect of usFJ *vs*. 6X for wild-type RNAP; Figure 7B). Given that βR542 interacts with the dsFJ, these data suggest that βR542 contributes to elemental pausing but less on a mutant template altered near its contact. We conclude that βR542 may help inhibit template dC loading in the ePEC.

**FIGURE 7.**
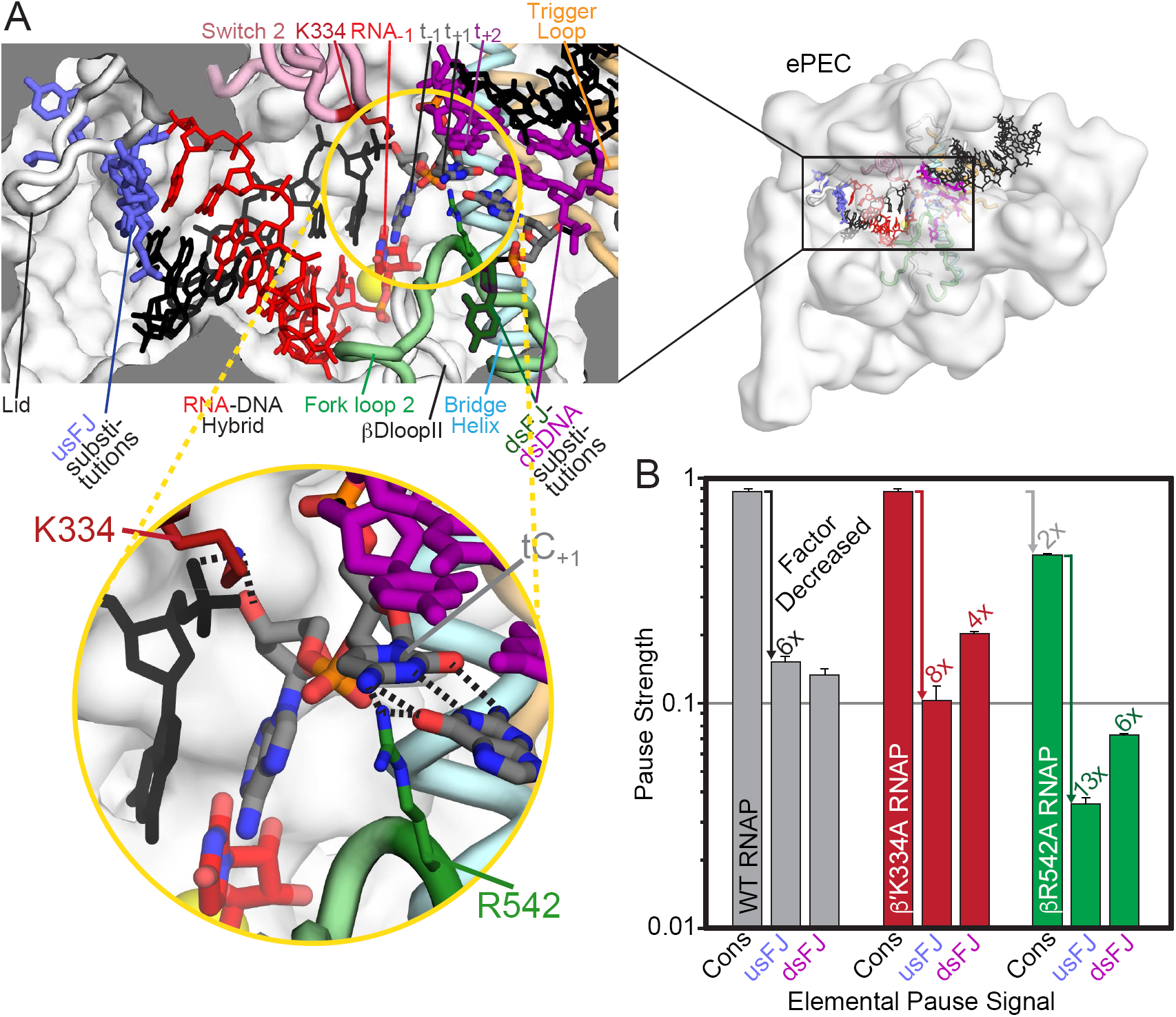
β R542 in fork-loop 2 may contribute to template base loading barrier in ePEC. Locations of β’K334 and βR542 in the ePEC. Relevant components of the ePEC are colored and labeled in a cutaway view of the active-site region of the ePEC. (**B**) Relative pause strength (PS_P_, Figure 2B) for wild type, β’K334A, and βR542A RNAPs on the consensus pause scaffold (Cons), the usFJ mutant scaffold, and the dsFJ mutant scaffold (see Figure 2A). The factor decrease relative to the wild-type RNAP on the consensus pause scaffold or relative to the consensus pause for the different RNAPs is given above the bars and depicted using colored arrows. Error is s.d. from ≥ 3 replicates.

## DISCUSSION

Our mechanistic study of elemental pausing documents key features of this fundamental regulatory behavior of RNAP. The elemental pause is an offline state that forms in competition with rapid elongation; thus, the pause mechanism requires distinct EC and pause conformations rather than a single state with an energetic barrier to translocation. The elemental pause signal is multipartite. RNAP interactions with the usFJ, Hyb, dsFJ, and dsDNA contribute to the key barrier to pause escape: template-base loading from a half-translocated state that involves clamp loosening and βR542 interactions. Other translocation states that can prolong ePEC lifetime form readily. Based on these results, we will describe a model for ePEC formation and escape and discuss its implications for gene regulation.

### A model for elemental pause formation and escape

To explain how RNAP responds to an elemental pause signal, we propose a multi-state pause mechanism in which pause escape is principally inhibited by inability to load the template base into the active site of RNAP in a half-translocated, off-line intermediate (Figure 8A,B). When RNAP encounters an elemental pause signal, a modest shift in the mobile modules of RNAP that are in contact with RNA and DNA (the clamp, shelf, lid, rudder, switch regions, lobe, protrusion, fork loop 2, and βDloopII) occurs during translocation. This shift creates or reinforces an energetic barrier to completion of translocation from a half-translocated state in which the RNA transcript has translocated but the DNA template has not, corresponding to the tilted hybrid intermediate observed in cryo-EM structures (Kang *et al.*, 2018a; Guo *et al.*, 2018). Comparison of ePEC and posttranslocated EC cryoEM structures reveals movements that slightly reposition these key mobile modules (compare green EC to magenta ePEC positions, Figure 8A). Key contacts preventing DNA translocation involve the lid, rudder, and switch 2 (Kang *et al.*, 2018a; Guo *et al.*, 2018) as well as apparent H-bonds of R542 in fork loop 2 to the template dC that may hinder its translocation into the active site (Figures 7 and 8A; compare to the conserved Arg in a *Tth*EC, which contacts the backbone phosphate of a template nucleotide loaded into the active site; Vassylyev *et al.*, 2007). The fraction of EC that partitions into the ePEC state and the height of the energetic barrier to completion of translocation and pause escape are both functions of the specific sequences present in the elemental pause signal (*i.e.*, a suboptimal signal will capture fewer ECs for a shorter overall dwell time).

**FIGURE 8.**
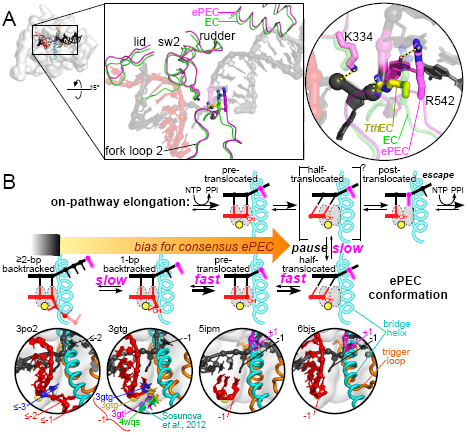
Multistate model of elemental pausing. (**A**) The small shift in RNAP modules observed in the ePEC (pdb 6bjs; magenta) relative to an EC (pdb 6alf; green) is depicted in the top central panel (Kang *et al.*, 2018a; Kang *et al.*, 2017). The locations of K334 in the EC (faint green) and ePEC (magenta) and the locations of R542 in a *Tth*RNAP EC (pdb 2o5i, yellow; Vassylyev *et al.*, 2007), EC, and ePEC are shown in the right panel expanded view. The +1 dC nucleotide trapped in the downstream DNA is magenta. (**B**) Discrete states present during formation of and escape from an elemental pause. The kinetic block is the loading of the template base, but the ePEC can assume other translocation registers depending on the sequences surrounding the pause. Examples of structures occupying the different translocation registers are shown in the close-ups below the active-site schematics.

DNA and RNA in the ePEC can move backwards to form pretranslocated, frayed, 1-bp backtracked, and, much more slowly, ≥2-bp backtracked states (Figure 8B). At least at the consensus pause, the half-translocated, pretranslocated, and 1-bp backtracked states equilibrate but these equilibria are biased toward the half-translocated state. This bias explains why we observed a fast translocation signal for the RNA:DNA hybrid (Figure 3) and the half-translocated intermediate in reconstituted ePECs by cryo-EM (Kang *et al.*, 2018a; Guo *et al.*, 2018). The fast equilibria explain why GreA rapidly cleaves a 1-bp backtracked ePEC state (Figures 1H&I; Figure S3); the 1-bp backtracked state quickly repopulates after cleavage even though it is only a small fraction of the total ePEC states. It is possible that GreA shifts the bias toward the 1-bp backtracked state, but mass action “pulling” of ePECs into the 1-bp backtracked state is a sufficient explanation. Because the cleaved ePEC re-encounters a strong pause signal, we observed little effect of GreA cleavage on pause lifetime. Our Gre A/B cleavage data also suggest a much slower entry into ≥2-bp backtracked states, which may be related to the minor but variable slow fraction of ePECs (see below).

This multistate model of the elemental pause predicts that the pretranslocated and 1-bp backtracked states can contribute significantly to pause lifetime at some pause sequences despite being in rapid equilibrium with the half-translocated intermediate in which the kinetic barrier to pause escape (template-base loading) is manifest. If an ePEC spends only 50% of the time half-translocated and 50% pre-translocated or backtracked, then the pause dwell time will increase by a factor of ∼2 relative to an ePEC that spends >95% of the time half-translocated. This is true despite the rapid equilibria because only the half-translocated intermediate can escape the pause and the other states diminish its effective concentration. Shifting to 25% or 10% half-translocated would lengthen the pause by factors of ∼3 or ∼9, respectively, for the same reason. Thus, an elemental pause signal that favors the 1-bp backtracked state, although different from the consensus signal we studied, would increase pause lifetime. These behaviors have been observed *in vitro* directly (Gabizon *et al.*, 2018) or by biasing ePECs using opposing force (Galburt *et al.*, 2007; Dangkulwanich *et al.*, 2013), as well as *in vivo* (Imashimizu *et al.*, 2015). Further, multiple locations of the 1-bp backtracked RNA 3’ nucleotide have been observed or proposed (Figure 8B; Sekine *et al.*, 2015; Wang *et al.*, 2009; Sosunova *et al.*, 2013). From a mechanistic standpoint, possible entry into multiple 1-bp backtracked states means that their contribution to pause dwell times may be both increased and highly variable as a function of sequence. The multistate elemental pause model makes the distinction between backtrack pausing and non-backtrack pausing a matter of degree rather than a clear-cut division.

Our results do not address the important question of whether the half-translocated state also is a significant kinetic intermediate during on-pathway nucleotide addition, although other studies suggest this possibility. Translocation can affect overall elongation rate (Imashimizu *et al.*, 2013; Gabizon *et al.*, 2018), and a half-translocated intermediate has been directly observed during *in crystallo* nucleotide addition by RNA-dependent RNAP (Shu and Gong, 2016). On-pathway translocation may proceed *via* initial RNA translocation then template DNA loading; the second step may be naturally slower at a pause signal, allowing time for formation of the ePEC conformation in which template-base loading becomes strongly inhibited (Gabizon *et al.*, 2018).

The multistate model (Figure 8) has important regulatory implications. Just as different regulators can exert distinct effects on the multistep mechanism of transcription initiation by stabilizing or destabilizing different intermediates (Hubin *et al.*, 2017), the existence of multiple elemental pause states affords multiple targets for regulators even when these intermediates are in equilibrium. For example, ppGpp is known to stimulate pausing (Kingston *et al.*, 1981) and to promote backtracking (Kamarthapu *et al.*, 2016); ppGpp could stimulate pausing by stabilizing a pretranslocated or backtracked PEC. Further, ≥1-bp backtracking may become more significant for other sequence contexts, conditions, or RNAPs (*e.g.*, eukaryotic RNAPII).

### Multipartite RNAP contacts, not RNA-DNA energetics alone, drive elemental pausing

Our results (Figure 2) confirm that, like the hairpin-stabilized *his*PEC (Chan *et al.*, 1997), the elemental pause signal is multipartite and involves significant contributions from the usFJ, hybrid, dsFJ, and dsDNA. Although strong evidence of significant contributions by the hybrid and dsDNA exists (Bochkareva *et al.*, 2012; Larson *et al.*, 2014; Palangat and Landick, 2001; Palangat *et al.*, 2004), some descriptions of pausing focus only on the usFJ and dsFJ (Vvedenskaya *et al.*, 2014; Imashimizu *et al.*, 2015). The inhibitory contributions of the usFJ and dsFJ to hybrid unwinding and NTP binding, respectively, have been known since the earliest studies of pausing (Gilbert *et al.*, 1974; Aivazashvili *et al.*, 1981) but alone are inadequate to predict pause strength.

At least three factors may contribute to underestimation of the importance of hybrid and downstream DNA sequences in pausing. First, the widespread use of sequence logos to represent nucleic acid signals places undue weight on simple, independent interactions and underweights contributions of complex sequence interactions that affect energetics through nucleic-acid conformation or alternative side-chain contacts. Thus, a logo representing information content as single bp can cause complex, multipartite sequences to appear less important (Figure 2A). Although more sophisticated algorithms hold promise to characterize complex, multipartite signals like the elemental pause (Siebert and Söding, 2016), direct analyses of sequence variants remain the most reliable way to define relevant contributions. Direct mutational analyses establish the key contributions of the hybrid and downstream DNA in the elemental pause signal (Figure 2; Bochkareva *et al.*, 2012; Larson *et al.*, 2014; Palangat and Landick, 2001; Palangat *et al.*, 2004).

Second, using NET-seq to identify a consensus pause signal requires rapid capture of PECs by quick-freezing actively growing cells in liquid N_2_ (Churchman and Weissman, 2012; Larson *et al.*, 2014). Quick-freezing reveals the modest sequence signatures of the hybrid and dsDNA in sequence logos and energetic analyses (Figure 2A; Larson *et al.*, 2014). However, some studies that detected little if any hybrid or dsDNA contribution recovered cells by centrifugation before freezing, which may allow RNAP to escape all but the strongest pause sites and thus could over-represent contributions of the usFJ and dsFJ components (Vvedenskaya *et al.*, 2014; Imashimizu *et al.*, 2015).

Finally, the hybrid and dsDNA contact multiple RNAP modules (*e.g*., rudder, switch 2, clamp, *etc*.) and may principally affect the complex and modest conformational rearrangement into the ePEC state. Because this conformation is still incompletely understood, the contributions of the hybrid and dsDNA, which may involve specific duplex conformations, may be difficult to characterize.

Our findings taken together with earlier demonstrations of usFJ, hybrid, dsFJ, and dsDNA sequences that contribute to pausing should solidify a model of elemental pausing that depends on a multipartite pause signal.

### Structure of the elemental pause

The elemental pause is a distinct offline state, not an on-pathway elongation intermediate. Our study uncovered evidence for a modestly rearranged multistate ePEC in which the clamp is loosened and template-base loading is inhibited as well as a slower ePEC state entered by a minor but variable fraction of ECs (Figures 1, 2, and S4). The lack of effect of rapid GreA cleavage ruled out 1-bp backtracking as an explanation of the slow ePEC state, but the GreA/B experiments could not definitively rule out ≥2-bp backtracking because the maximal ≥3-nt cleavage rate of ∼0.002 s^-1^ (Zhang *et al.*, 2010; Sosunova *et al.*, 2013) is too close to slow ePEC escape rate. Alternatively or additionally, the slow ePEC state could involve RNAP swiveling, which is observed in the hairpin-stabilized PEC (Kang *et al.*, 2018b). Swiveling involves a near-rigid body rotation of the clamp, shelf, SI3, and jaw, and inhibits nucleotide addition by a factor of ∼10 by interfering with TL folding. Interestingly, the lifetime of the minor slow ePEC is ∼10 times that of the majority ePEC state. Variations in susceptibility to swiveling, ≥2-bp backtracking, or both could explain why the slow fraction varies among RNAP preparations. Further studies will be needed to establish the structure and importance of the slow ePEC fraction, including detection with non-reconstituted transcription complexes or *in vivo*.

The multistate model of elemental pausing involving small changes in RNAP conformation and half-translocated, pretranslocated, and multiple 1-bp backtracked states described here will be difficult to test fully using ensemble biochemistry, single-molecule biochemistry, or crystallography. These methods are encumbered by the perturbing effects of probes and experimental configurations, the number of intermediates involved, and the rapid time-scale of their interchange. Time-resolved cryo-EM (Frank, 2017) promises an attractive approach if methods to distinguish intermediates during particle classification can be developed.

## MATERIALS AND METHODS

### Reagents and materials

Plasmids and oligonucleotides are listed in Table S1. RNA and DNA oligonucleotides were obtained from Integrated DNA Technologies (IDT; Coralville, IA) and purified by denaturing polyacrylamide gel electrophoresis (PAGE) before use. [γ-^32^P]ATP, [α-^32^P]CTP and [α-^32^P]GTP were obtained from PerkinElmer Life Sciences; rNTPs, from Promega (Madison, WI); and 6-MI, from Fidelity Systems (Gaitersburg, MD). RNAPs were purified as described previously (Windgassen *et al.*, 2014). Briefly, His-tagged proteins were overexpressed in *E. coli* BL21 (DE3) and cells were lysed by sonication. RNAPs were enriched by PEI and ammonium sulfate precipitation, then purified by sequential nickel (5 mL HisTrap) and heparin (5 mL HiTrap) column chromatography, dialyzed into storage buffer (20 mM Tris-Cl, pH 8, 250 mM NaCl, 20 μM ZnCl_2_, 1 mM MgCl_2_, 0.1 mM EDTA, 1 mM DTT, and 25% glycerol), and stored in small aliquots at –80° C.

### *In vitro* transcription pause assays

PAGE-purified 15-mer RNA with 3’ end 2 nt upstream from the pause site (5 µM) and template DNA (10 µM) were annealed in transcription buffer 1 (TB1; 20 mM Tris-OAc pH 7.7, 5 mM Mg(OAc)_2_, 40 mM KOAc, 1 mM DTT). The sequences of the nucleic acids and their corresponding stock numbers are listed in the Key Resources Table. Scaffolds were incubated with RNAP for 15 min at 37 °C in TB1, then non-template DNA was added and incubation continued for 15 min at 37 °C. The ratio of RNA:tDNA:RNAP:ntDNA was 1:2:3:5 (0.5 µM, 1 µM, 1.5 µM, 2.5 µM, respectively). ECs were diluted to 0.1 µM with TB1 + heparin (0.1 mg/ml), incubated for 3 min at 37 °C, labeled by the incorporation of [α-^32^P]GMP at 10 µM total GTP for 1 min at 37 °C, and then placed on ice for 30-60 min. ECs were incubated for 3 min at 37 °C before initiating the pause assay by addition of CTP to 100 µM and GTP to 10 or 100 µM in TB1 at 37 °C. Reaction samples were removed at various time points and quenched with an equal volume of 2X stop buffer (8 M urea, 50 mM EDTA, 90 mM Tris-borate buffer, pH 8.3, 0.02% each bromophenol blue and xylene cyanol). All remaining active ECs were chased to product by incubation with GTP at 1 mM for 1 min at 37 °C. RNAs in each quenched reaction sample were separated on a 15% PAG (19:1 acrylamide:bis-acrylamide) in 44 mM Tris-borate, pH 8.3, 1.25 mM Na_2_EDTA, 8 M urea. The gel was exposed to a PhosphorImager screen, and the screen was scanned using Typhoon PhosphorImager software and quantified in ImageQuant. The averaged fraction of RNA at the position of the pause over time was fitted to single- or double-exponential decay functions in KaleidaGraph to estimate pause efficiencies (amplitudes) and rate constants of pause escape. For experiments comparing RNAPs, wild-type and variant RNAPs were purified side-by-side to avoid variable effects of different RNAP preparations on pausing kinetics.

### Kinetic modeling

To test whether the elemental pause is on online or offline state (*i.e*., involves a linear or branched kinetic mechanism; Figures S1F and S1G) and to test whether the slow fraction of ePECs was evident at only the first or at both pause sites on the tandem pause scaffold (Figure S2), we used kinetic modeling by numerical integration of pre-steady state rate equations using the program KinTek Explorer v6.1 (KinTek Corp., Snow Shoe, PA; Johnson *et al.*, 2009). In both cases, to test whether the simpler mechanism was adequate to explain the data, we used replicate datasets (triplicate or greater) to generate a kinetic model for the rates of arrival at the pause site using the rate at which all RNAs before the pause site converted to RNAs at the pause site and beyond. We then held these rates constant and tested the simple kinetic models (linear, online pause (Figures S1F and S1G) or two populations of RNAP, fast and slow pausing (Figures S2D and S2E), including error for replicates to obtain the best fit and the residuals between the best fit and the observed data (see Figures S1F,G and S2D,E). We concluded that the more complex models (branched for Figures S1F,G or dynamic formation of the slow pause species for Figures S2D,E) were favored because the residuals for the simpler mechanism exhibited large, systematic variations whereas the more complex mechanisms exhibited smaller, random variations. We did not attempt to determine which mechanism best fit the data, and limited our conclusion to rejection of the simpler mechanism. For the dataset using the template from Bochkareva et al., 2012 (Figure S1G), for which we had six replicates, we determined error in the fits and residuals by individually fitting each dataset and calculating the average and error for the six fits.

### Intrinsic cleavage assay

Intrinsic cleavage was assayed essentially as described earlier (Mishanina *et al.*, 2017). Briefly, 3’-end labeled C17 ePECs (#9563 NT DNA, #8334 T DNA, #8401 RNA; Table S1) were formed and immobilized on Ni^2+^-NTA beads, then washed to remove unincorporated [α-^32^P]CTP. Cleavage was initiated at 37 °C by resuspending washed ePECs with Cleavage Buffer (CB; 25 mM Tris·HCl pH 9.0, 50 mM KCl, 20 mM MgCl_2_, 1 mM DTT, 5% glycerol, and 25 µg acetylated BSA/mL), and samples were collected at designated timepoints by mixing with 2X stop buffer. Cleavage products were separated by denaturing PAGE as described for transcription pause assays.

### Stopped-flow fluorescence translocation assay

To measure translocation rates of the hybrid (Figure 3), we used the assay developed by Belogurov and co-workers (Malinen *et al.*, 2012; Malinen *et al.*, 2014). PAGE-purified template DNA and RNA were annealed in TB1 (see Table S1). This scaffold was incubated with RNAP for 15 min at 37 °C in TB, then non-template DNA was added such that the final ratio of tDNA:RNA:RNAP:ntDNA was 1:2:3:5 (2 µM: 4 µM: 6 µM: 10 µM, respectively). This solution was diluted to 0.4 µM RNAP with TB1.

ECs were then injected into one loading syringe of a stopped-flow apparatus (SF-300X; KinTek Corporation, Snow Shoe, PA) and 200 µM CTP in TB1 was loaded in the other syringe. Upon initiating rapid mixing at 37 °C, 6-MI fluorescence was excited at 340 nm (2.4 nm bandwidth), and emission was monitored in real time through a 400 nm long-pass filter (Edmund Optics Inc., Barrington, NJ). The kinetics of 6-MI fluorescence unquenching was determined by fitting the average fluorescence (n ≥6 traces), normalized from 0 to 1, to a double exponential equation [1]:

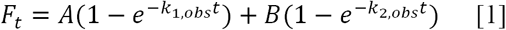

where, t = time (s); A = fast kinetic species signal amplitude; B = slow kinetic species signal amplitude; *k*_1,obs_ = observed rate of fluorescence increase for the fast kinetic species; *k*_*2,obs*_ = observed rate of fluorescence increase for the slow kinetic species. The fast species rate is reported.

### Rapid quench-flow measurement of nucleotide addition

To measure rates of C17 and G18 addition, G16 ECs were formed essentially as described for the pause transcription assay, but with 5’-[^32^P]RNA limiting such that RNA:tDNA:RNAP:ntDNA was 1:1.3:2:3.3. To obtain nucleotide addition rates using a quench-flow apparatus (RQF-3; KinTek Corporation, Snow Shoe, PA), 400 nM G16 ECs were injected in one sample loop and 200 µM each CTP and GTP in TB1, in the other sample loop. Reactions were performed at 37° C for the designated times and quenched with 2 M HCl, then neutralized immediately to pH 7.8 with 3 M Tris base (supplemented with 250 µg torula yeast RNA/mL). RNA products were purified by phenol:chloroform extraction followed by ethanol precipitation, and resuspended in 1X stop buffer to a constant specific activity. RNA products from all timepoints were resolved by denaturing PAGE and quantified as described for pause transcription assays.

Reaction progress curves were generated for each RNA length (G16, C17^+^, and G18^+^) using KaleidaGraph (Synergy Software) by calculating the fraction of total RNA for each condition as a function of time. C17 and all RNAs longer than C17 were combined to give the C17^+^ fraction; G18 and all RNAs longer than G18 were combined to give the G18^+^ fraction. The averaged fraction at each time-point was then fit to a single-exponential equation [2]:

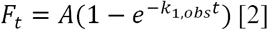

### Equilibrium fluorescence measurement of translocation

To measure equilibrium translocation of downstream DNA (Figure 4), G16 ECs were formed in TB1 as described for the quench-flow experiment but without Mg(OAc)_2_ to stabilize ECs and with the fluorescent ntDNA as the limiting component (ntDNA:tDNA:RNA:RNAP = 1:1.5:2:3 with ntDNA at 300 nM). For equilibrium measurements of hybrid translocation (Figure 3), G16 ECs were formed similarly but with the tDNA as the limiting component (see description in stopped-flow assay section). Fluorescence measurements were conducted using a PTI-spectrofluorometer (Model QM-4/2003, Photon Technology International) with 5-mM path length and 45-uL quartz cuvettes (Hellma Analytics, Müllheim, Germany). Emission spectra were obtained by exciting at 340 nm (5 nm bandwidth) and monitoring fluorescence between 360 and 500 nm (5-nm bandwidth).

For each substrate addition experiment, 60 µL EC was added to the cuvette, incubated for 2 min at 37 °C in the cuvette holder before performing an emission scan (average of 3 traces at 0.25 nm step size). Then, NTPs were added (concurrently with 5 mM Mg(OAc)_2_ plus additional Mg(OAc)_2_ equal to the NTP concentration in the assay) and the incubation was continued for 1 min (5 min for 2’dCTP) at 37 °C before performing an emission scan. Between fluorescence measurements, an aliquot was removed to 2X stop buffer to run on denaturing PAGE to confirm nucleotide addition. For the GTP titration condition, aliquots were only removed to confirm initial G16 RNA, 3’dC addition, and failure of 20 mM GTP to incorporate following 3’dC addition.

6-MI fluorescence was quantified at 425 nm. Background fluorescence from a GTP contaminant was subtracted using signal from GTP alone in buffer. We found that the level of fluorescence from the contaminant varied significantly among vendors and GTP lot, with GTP from GE Healthcare containing the least amount. Reported increases in fluorescence are fold changes relative to the initial signal in G16 EC.

### Cys-pair reporters (CPR) crosslinking assays

CPR crosslinking assays (Figure 5) were performed as described previously (Nayak *et al.*, 2013). Nucleic acid scaffolds were prepared by annealing RNA, template DNA (T-DNA) and 15 μM non-template DNA (NT-DNA) at 10 µM, 12 µM, and 15 µM final concentrations, respectively, in reconstitution buffer (RB; 20 mM Tris-HCl pH 8, 20 mM NaCl and 1 mM EDTA). ECs, ePECs, and *his* PEC were formed by incubating 1 μM RNAP and scaffold (2 μM, based on RNA) in buffer A (50 mM Tris-HCl pH 8, 20 mM NaCl, 10 mM MgCl_2_, 1 mM EDTA and 2.5 ug acetylated bovine serum albumin/mL for 15 min at room temperature (RT). For crosslinking reactions with NTP, 3’deoxyECs formed by reaction with 3’dNTP were incubated for 15 min at RT with 0, 0.005, 0.01, 0.025, 0.05, 0.1, 0.25, 0.5, 1, 2.5, 5, and 10 mM GTP or ATP. Next, EC, ePEC, or his-PEC (final RNAP 0.8 uM and scaffold 1.6 uM) were incubated for 60 min with 2.5 mM CSSC and 0.05 mM DTT (E = –0.16) and stopped with 50 mM iodoacetamide. Samples were separated by native PAGE to verify reconstitution efficiency and by sodium dodecyl sulfate (SDS)-PAGE using 4-15% GE Healthcare PhastGel to quantify formation of crosslinks. Gels were stained with Coomassie Blue and imaged with a CCD camera. The fraction cross-linked was quantified with ImageJ software. The experimental error was determined as the standard deviation of measurements from three or more independent replicates.

### Pause Assays with Cys-Pair Reporter RNAPs

Nucleic-acid scaffolds containing RNA and template DNA (1:2 ratio of RNA to DNA) were assembled for *in vitro* pause assays and used to reconstitute ePECs or control *his*PECs for Cys-pair crosslinking experiments (Figures 6C and 6D) as described in Kang *et al.*(2018a) with scaffolds indicated in the figure legends. The U15 ePECs containing limiting CPR RNAP (1 µM) were reconstituted on this scaffold (2 µM, based on RNA) for 15 min at 37 °C in Elongation Buffer (EB; 25 mM HEPES-KOH, pH 8.0, 130 mM KCl, 5 mM MgCl_2_, 1 mM DTT, 0.15 mM EDTA, 5% glycerol, and 25 µg acetylated bovine serum albumin/mL), followed by addition of 6 µM non-template DNA and further incubation for 10 min at 37 °C to complete assembly of the transcription complexes. Wild-type RNAP was tested as a control side-by-side with CPR RNAPs. Crosslinking of 1 µM ePECs was performed in the presence of 1 mM cystamine as the oxidant and 0.8 mM DTT, for 15 min at 37 °C. An aliquot of the crosslinking reaction was quenched with 15 mM iodoacetamide (final concentration) and analyzed by non-reducing SDS-PAGE to confirm formation of the crosslink.

The crosslinked U15 ePECs were diluted to 0.2 µM with EB (without DTT, for crosslinked samples) and incubated with heparin (0.1 mg/mL final) for 3 min at 37 °C. The U15 ePECs were then radiolabeled by extension with 20 µM [α-^32^P]GTP for 1 min at 37 °C, to poise the complexes one nucleotide before the pause sequence. The resulting G16 ePECs were further diluted to 0.1 µM (based on RNAP) and assayed at 37 °C for pause-escape kinetics at 10 µM GTP by addition of CTP in EB to 100 µM (without DTT, for crosslinked samples). Reaction samples were removed at various time points and quenched with an equal volume of 2X stop buffer. All active ePECs were chased out of the pause with 500 µM GTP and CTP, each, for 5 min at 37 °C. RNAs in each quenched reaction sample were separated on a 15% urea-PAGE gel. The gel was visualized and quantified as described for *in vitro* transcription assays.

### Cys Triplet Reporter Assays

Nucleic-acid scaffolds used to reconstitute hisPEC or ECs for Cys triplet reporter (CTR) cross-linking assays (Figures 6B, E and F) were assembled on purified DNA and RNA scaffolds shown in the figure legend as described previously (Kang *et al.*, 2018a). Briefly, 10 µM RNA, 12 µM template DNA, and 15 µM non-template DNA (Table S1) were annealed in RB. To assemble complexes, scaffold (2 µM) was mixed with limiting CTR RNAP (1 µM; CTR RNAP: β’1045iC 258iC, β843C) in 50 mM Tris-HCl, pH 7.9, 20 mM NaCl, 10 mM MgCl_2_, 0.1 mM EDTA, 5% glycerol, and 2.5 µg of acetylated bovine serum albumin/mL, and added to mixtures of cystamine and DTT to generate redox potentials that ranged from -0.314 to -0.424. Complexes were incubated for 60 min at room temperature and then were quenched with the addition of iodoacetamide to 15 mM. The formation of cysteine-pair cross-links was then evaluated by non-reducing SDS-PAGE (4%–15% gradient Phastgel; GE Healthcare) as described previously (Nayak *et al.*, 2013). Gels were stained with Coomassie Blue and imaged with a CCD camera. The fraction cross-linked was quantified with ImageJ software. The experimental error was determined as the standard deviation of measurements from three or more independent replicates.

## ACKNOWLEDGEMENTS

We thank members of the Landick lab for many helpful discussions during the course of this work and preparation of the manuscript. The work was supported by a grant from the NIH to R.L. (R01 GM38660).

## Supplemental Figure Legends

**TABLE S1.**
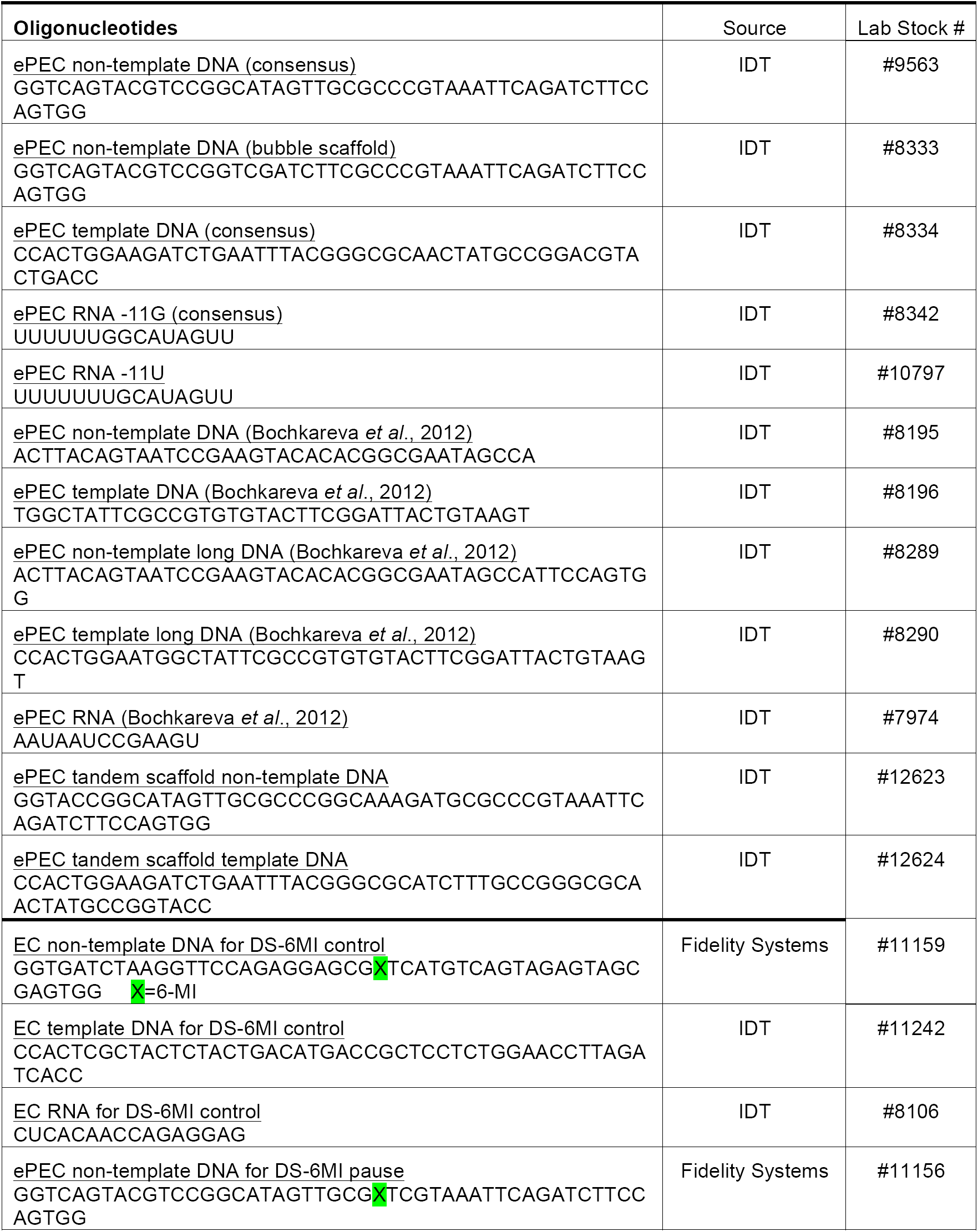
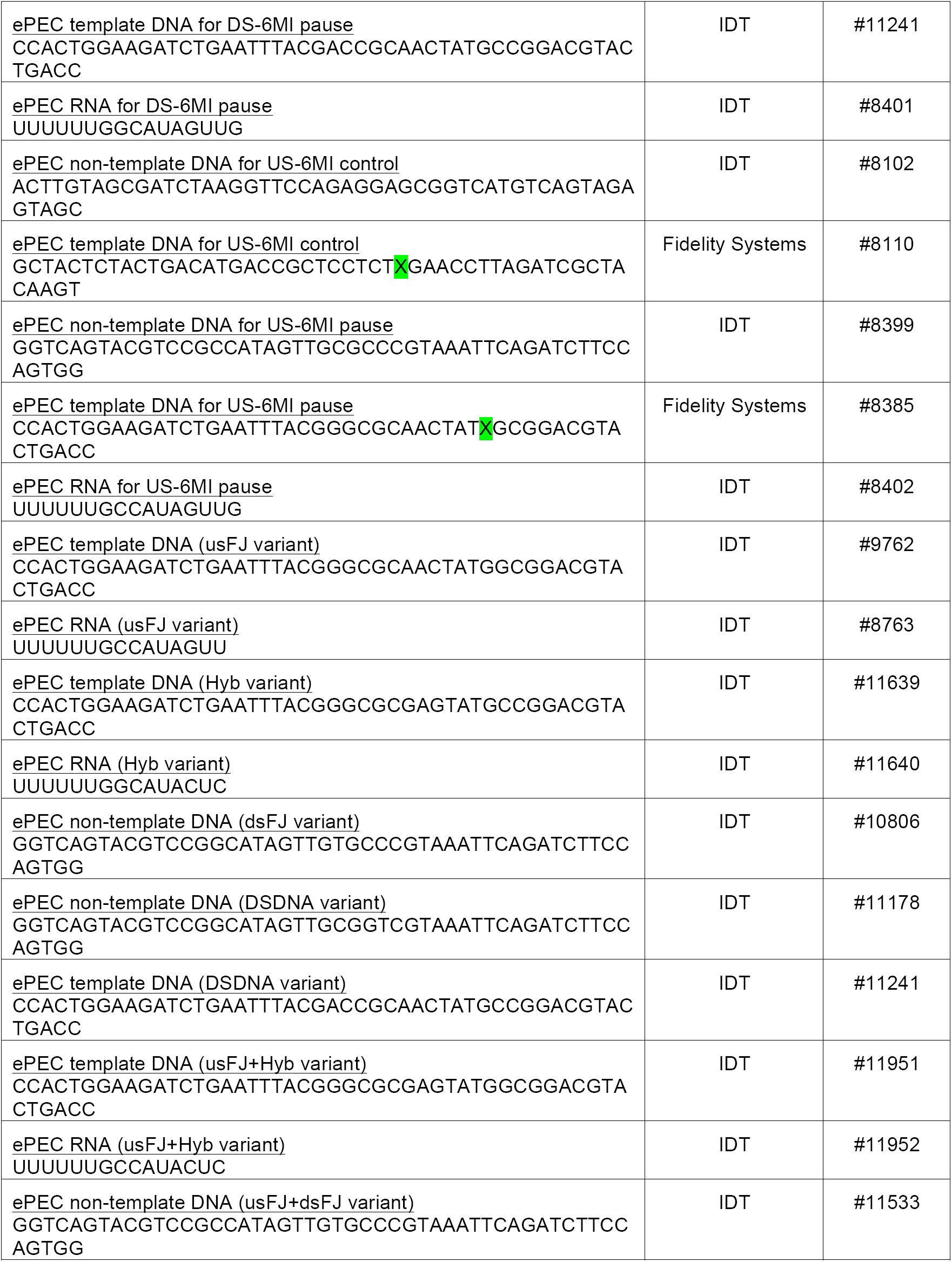
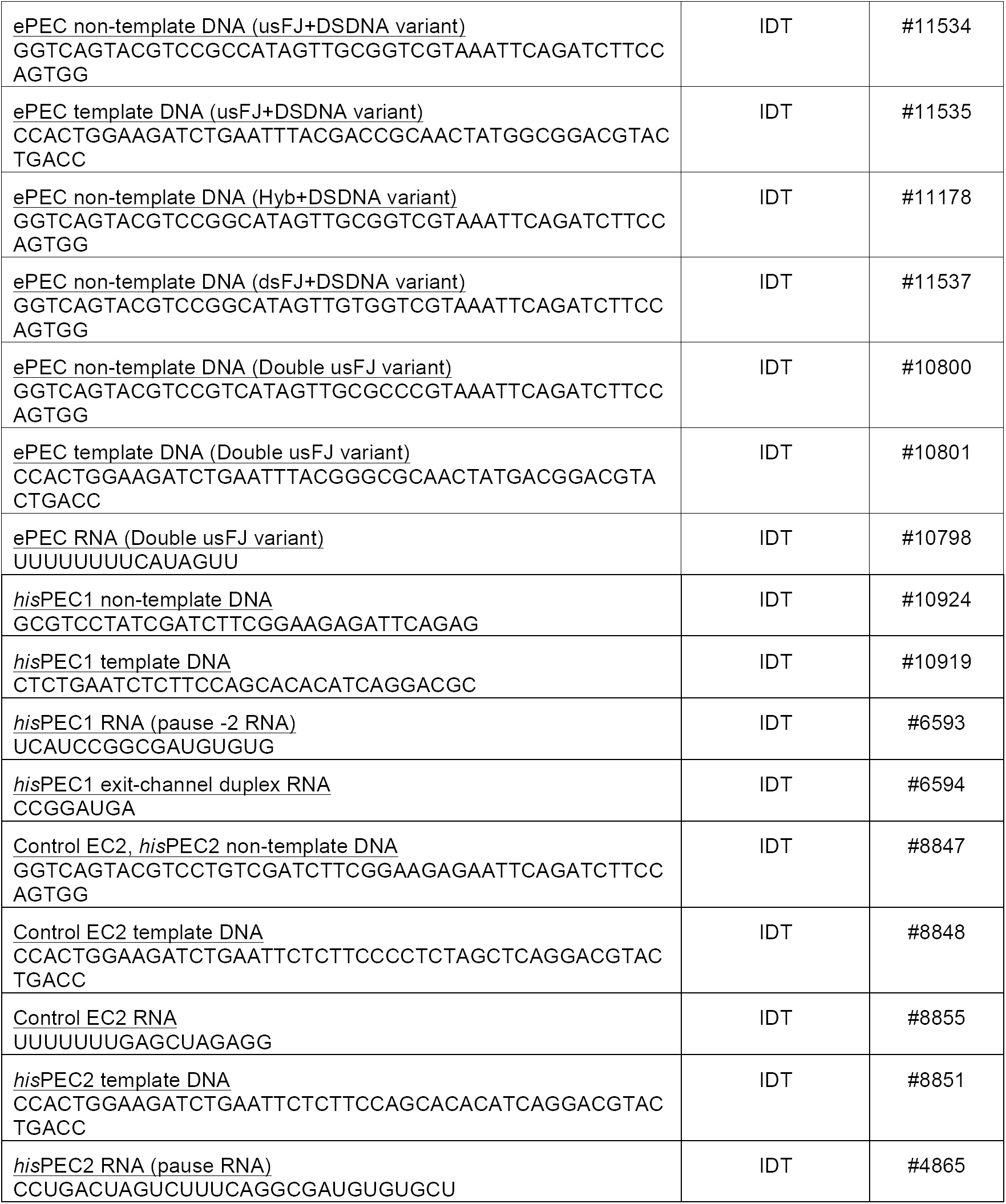
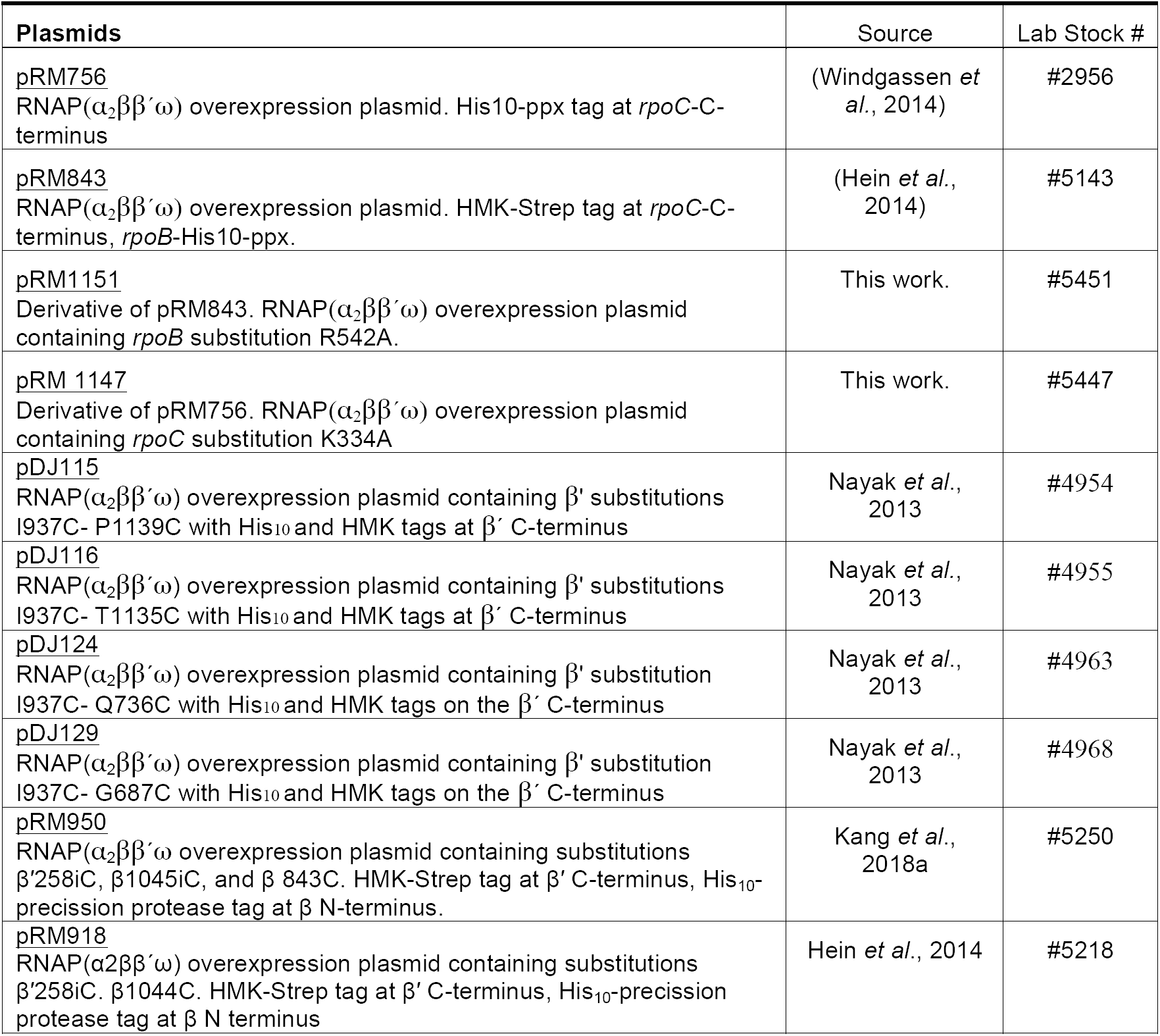

**Figure S1.**
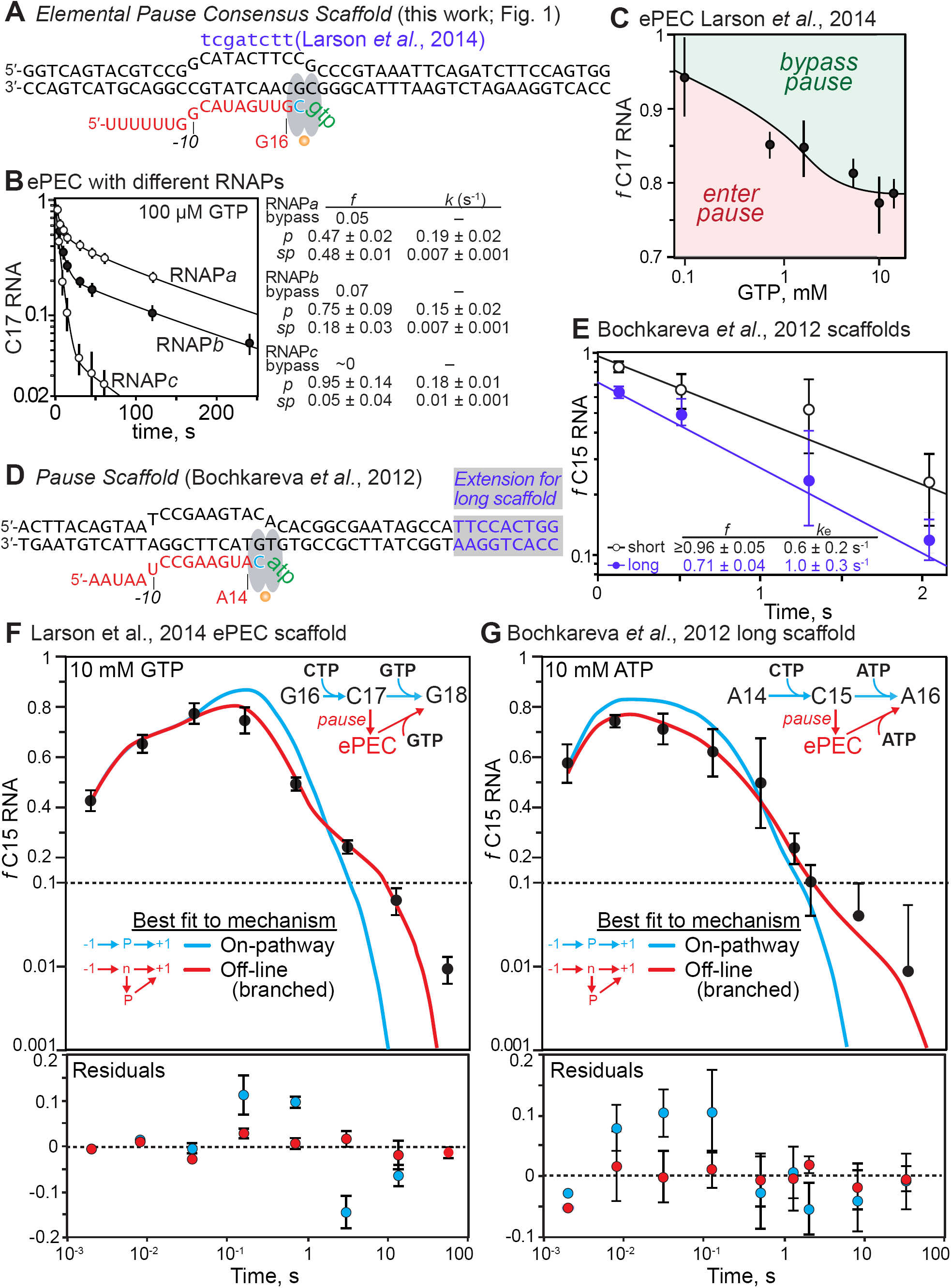
The elemental pause is a distinct offline state, not an on-pathway elongation intermediate. (**A**) The complete consensus elemental pause scaffold (ntDNA #9563, tDNA #8334, RNA #8342; Table S1), identical to that used in Larson *et al*., 2014, except the template and nontemplate DNA strands are fully complementary (the noncomplementary, nontemplate strand bases in the Larson *et al*. scaffold are indicated in purple; ntDNA #8333). (**B**) Pause RNA as a function of time and kinetic fractions for three different RNAP preparations. (**C**) Effect of GTP concentration on partition of C17 between bypass and pause states in experiments also reported in Larson et al., 2014. (**D**) The pause scaffold defined by Bochkareva et al., 2012 (*black*; ntDNA #8195, tDNA #8196, RNA, #7974; Table S1) and the 10 bp downstream duplex extension to move the duplex end outside the RNAP footprint (*purple* on *gray* background; *black*; ntDNA #8289, tDNA #8196). Steps in a minimal pause assay and the constrained offline pause pathway (*red*) are shown on the right. (**E**) Estimation of pause fractions and apparent pause escape rates for the Bochkareva *et al*. scaffold with and without 10 bp downstream duplex, extension at 10 mM ATP. (**F**) Best fit by numerical integration and residuals for on-pathway (unbranched) and off-line mechanisms for the consensus ePEC scaffold (panel A) at 10 mM GTP. The fits shown were calculated using the average of data from 3 experimental repeats, including error (s.d.) in the fitting. The residuals were calculating by performing individual fits to the 3 replicate datasets and then averaging and calculating s.d. for the residuals for each fit. The sinusoidally varying residuals for the unbranched mechanism *vs.* their random and within-error variation for the branched mechanism establishes that pausing occurs by the branched mechanism. (**G**) Best fit by numerical integration and residuals for on-pathway (unbranched) and off-line (distinct pause species at C15) mechanisms for the Bochkareva *et al.,* 2012 ePEC scaffold (panel B) at 10 mM ATP. The fits and residuals were calculated as described for panel E, except that data from 6 experimental repeats were used, and by the same reasoning establish that pausing occurs by the branched mechanism.

**Figure S2.**
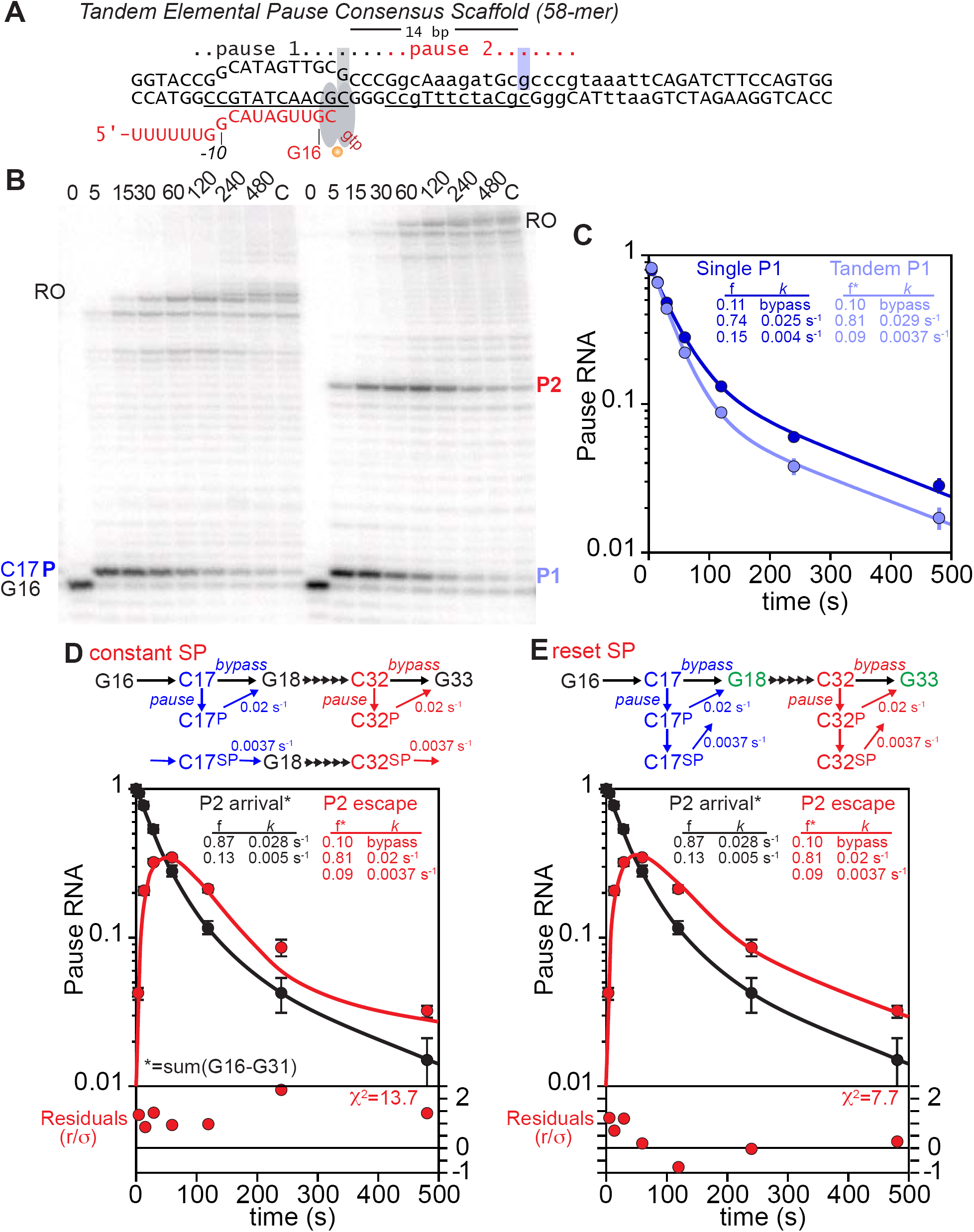
(**A**) Scaffold used to probe for biphasic escape kinetics at two sequential elemental pause sites (Tandem Pause Scaffold; tDNA #12623, ntDNA #12624, RNA #8342; Table S1). Pause assay with consensus pause scaffold (Single Pause; left time course) or Tandem Pause Scaffold (right time course). Transcription was initiated by addition of NTPs to 100 µM each ATP, UTP, and CTP, and 10 µM GTP. The gel image is representative of triplicate assays. Quantitation of the C17 pause band on the single or tandem pause scaffold. (**D**) Tandem scaffold pause P2 fit to a kinetic model with a single pause species. The fit was calculated by first modeling the arrival of ECs at the second pause, plotted as escape from the region before the pause (red line). This flux of ECs arriving at the second pause was subjected to least-squares fitting using numerical integration to a model of pausing involving only a single pause species. (**E**) Tandem scaffold pause P2 fit to a kinetic model with two pause species. Fitting was conducted as described for panel E, except the pause model included two pause species, with a slow pause arising from the faster pause.

**Figure S3.**
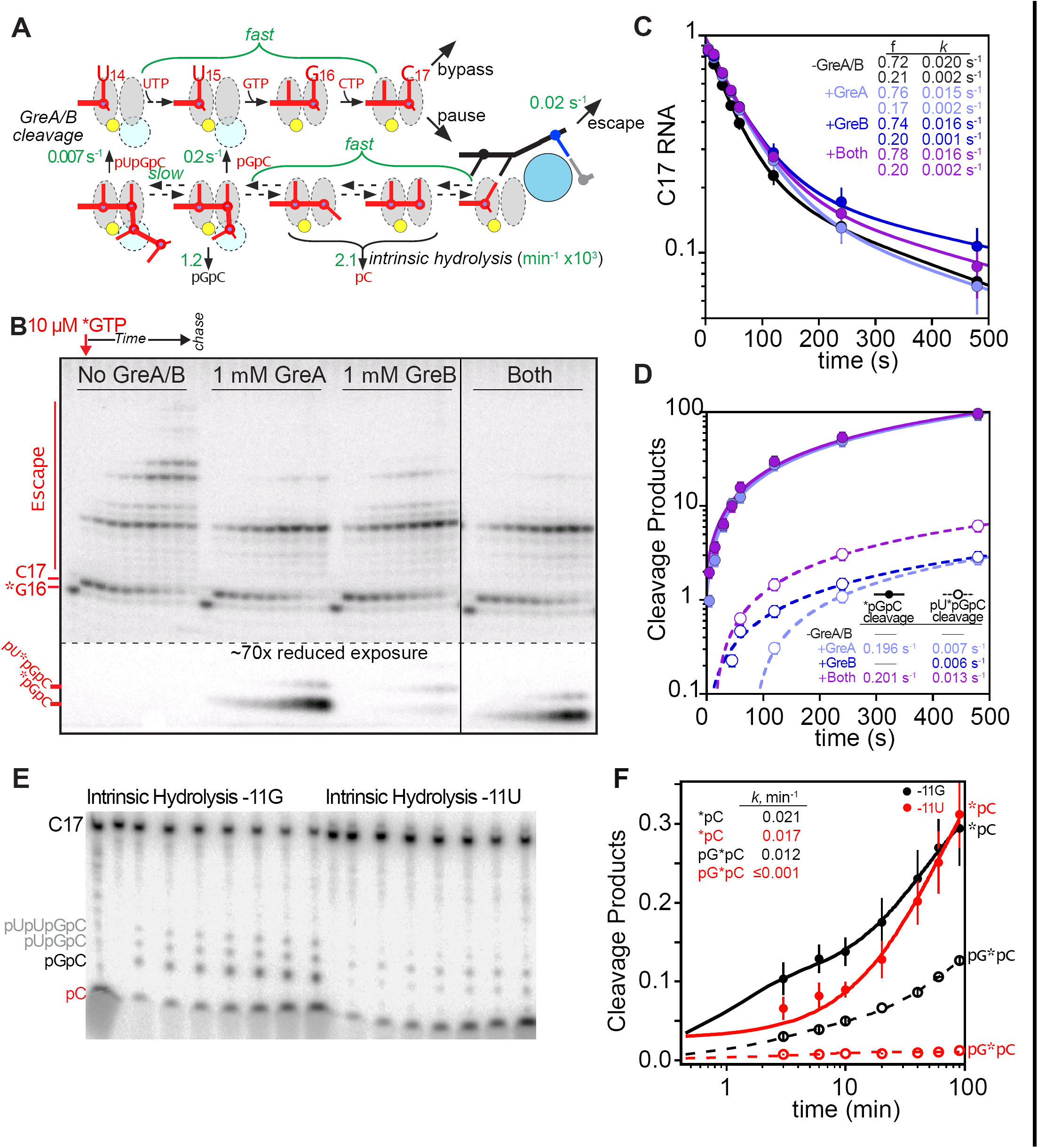
(**A**) Model for RNA cleavage in pre-translocated or backtracked states following entry into the elemental pause. Fast extension of ECs_U14-G16_ and fast interchange of half-translocated, pre-translocated, frayed, and 1-bp backtracked ePECs enables generate of 2-nt cleavage products even when the half-translocated ePEC in which the block to pause escape is manifest is the predominant species. Observed rates of GreA cleavage and intrinsic cleavage are indicated in green (from panels D and F). (**B**) Pause assays to test the effect of Gre factors. The RNA was 3’ end-labeled with 20 µM [α-^32^P]GMP (constant specific activity throughout the experiment). Transcription was initiated by addition of 2X NTPs (200 µM CTP, 200 µM UTP, – or + Gre factors). The gel image is representative of results obtained in at least triplicate assays. (**C**) Quantitation of pause fraction in B as a function of time. Error bars are SD of at least triplicates. (**D**) Quantitation of cleavage products in B as a function of time. Error bars are SD of at least triplicates. Cleavage rates too slow to measure (< 0.005 s^-1^) are indicated in the table as lines. (**E**) Intrinsic exo- and endonuclease activity on consensus pause scaffolds with -11G or -11U RNA (related to Figure 1J,K). After forming ECs and 3’-end labeling with [α-^32^P]CMP, the ECs were simultaneously used for pause assays (Figure 1K) and the intrinsic cleavage assays shown here. See methods for experimental details. (**F**) Quantitation of cleavage products in panel E as a function of time. The data were fitted to a double-exponential function. The rate constants for the major kinetic fractions are shown in the inset table.

**Figure S4.**
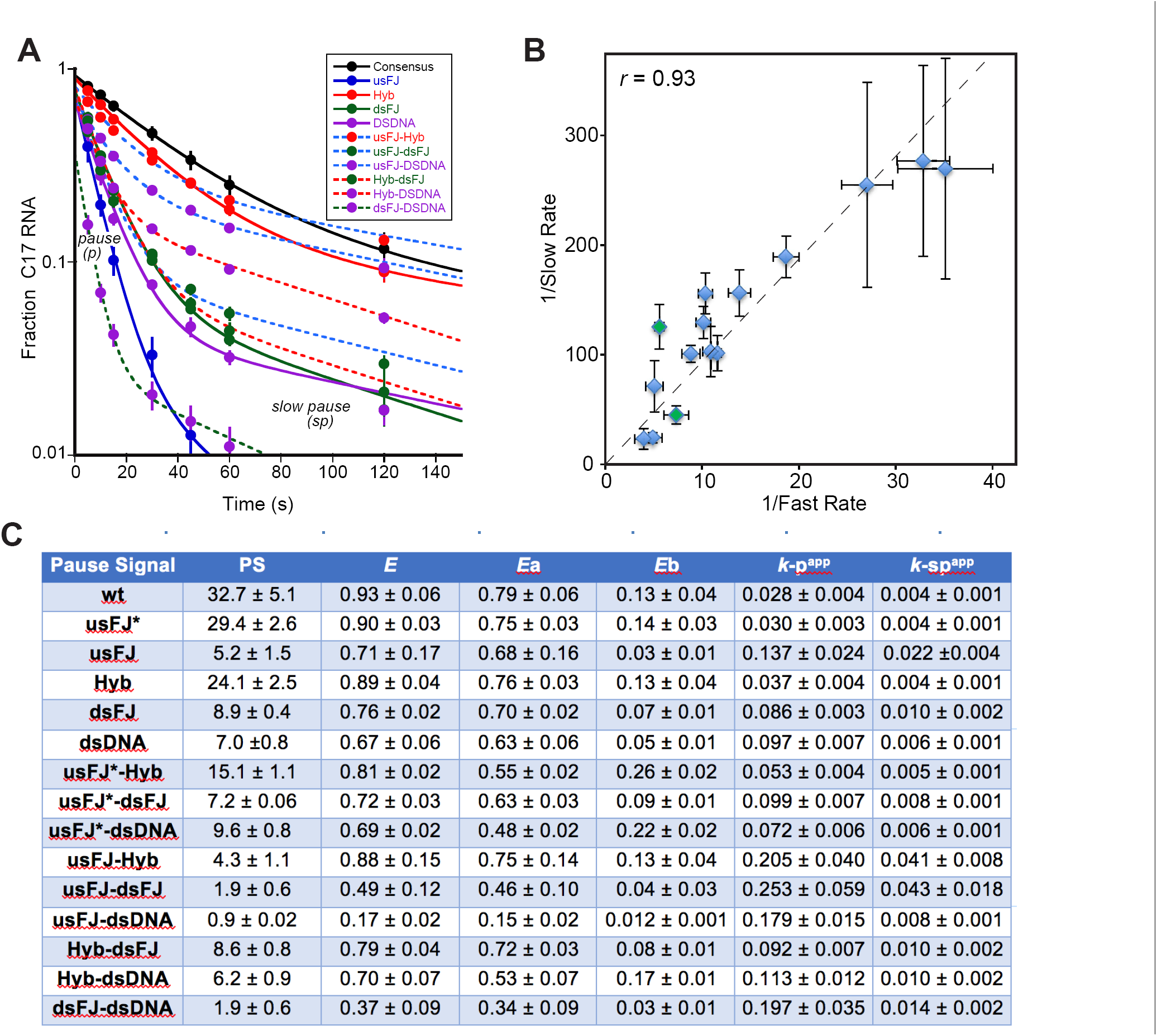
(**A**) Quantitation of pause assays with scaffold variants described in Figure 2. All pause assays were performed with the same RNAP preparation. (**B**) Plot of fast species rate (k_-p, app_) vs. slow rate (k_-sp, app_) for each scaffold variant. The high positive correlation between the two species rates is consistent with a model in which a single kinetic barrier limits escape from either pause species. (**C**) Summary table of pause strength (PS), efficiency (E), fast species fraction (*E*a), slow species fraction (*E*b), fast species rate (k_-p, app_), slow species rate (k_-sp, app_).

**Figure S5.**
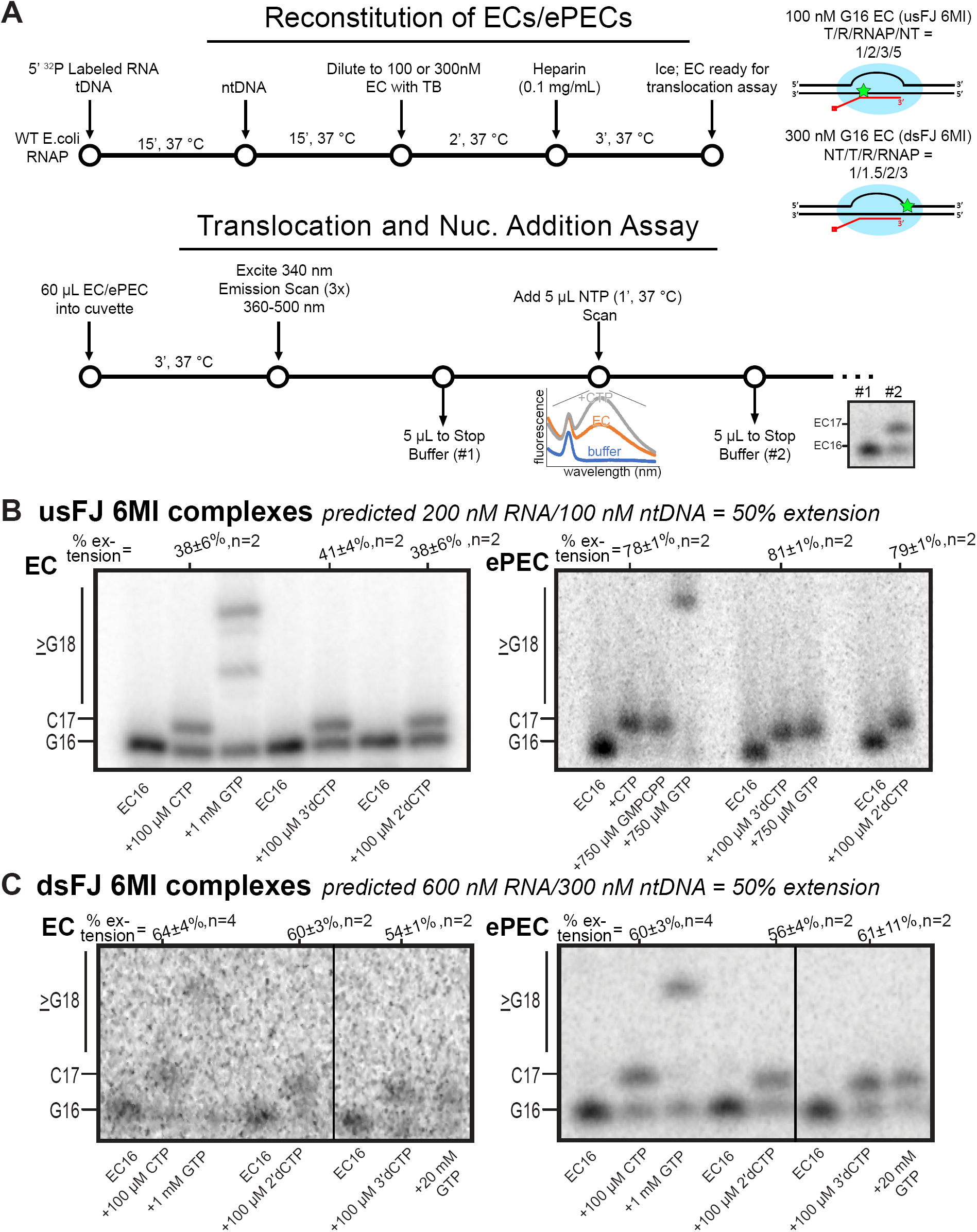
Reconstitution of ECs and PECs for 6-MI translocation assays. (**A**) Experimental schematic for the assay of translocation and nucleotide addition. (**B**) Nucleotide addition for samples used in translocation assays of usFJ 6MI complexes. Percentages above extension lanes indicate the fraction of RNA extended (vs. the prediction of 50% based on reconstitution ratios). Error is range for n=2 or s.d. for n≥3. (**C**) Nucleotide addition for samples used in translocation assays of dsFJ 6MI complexes. Percentages above extension lanes indicate the fraction of RNA extended (vs. the prediction of 50% based on reconstitution ratios). Error is range for n=2 or s.d. for n≥3.

**Figure S6.**
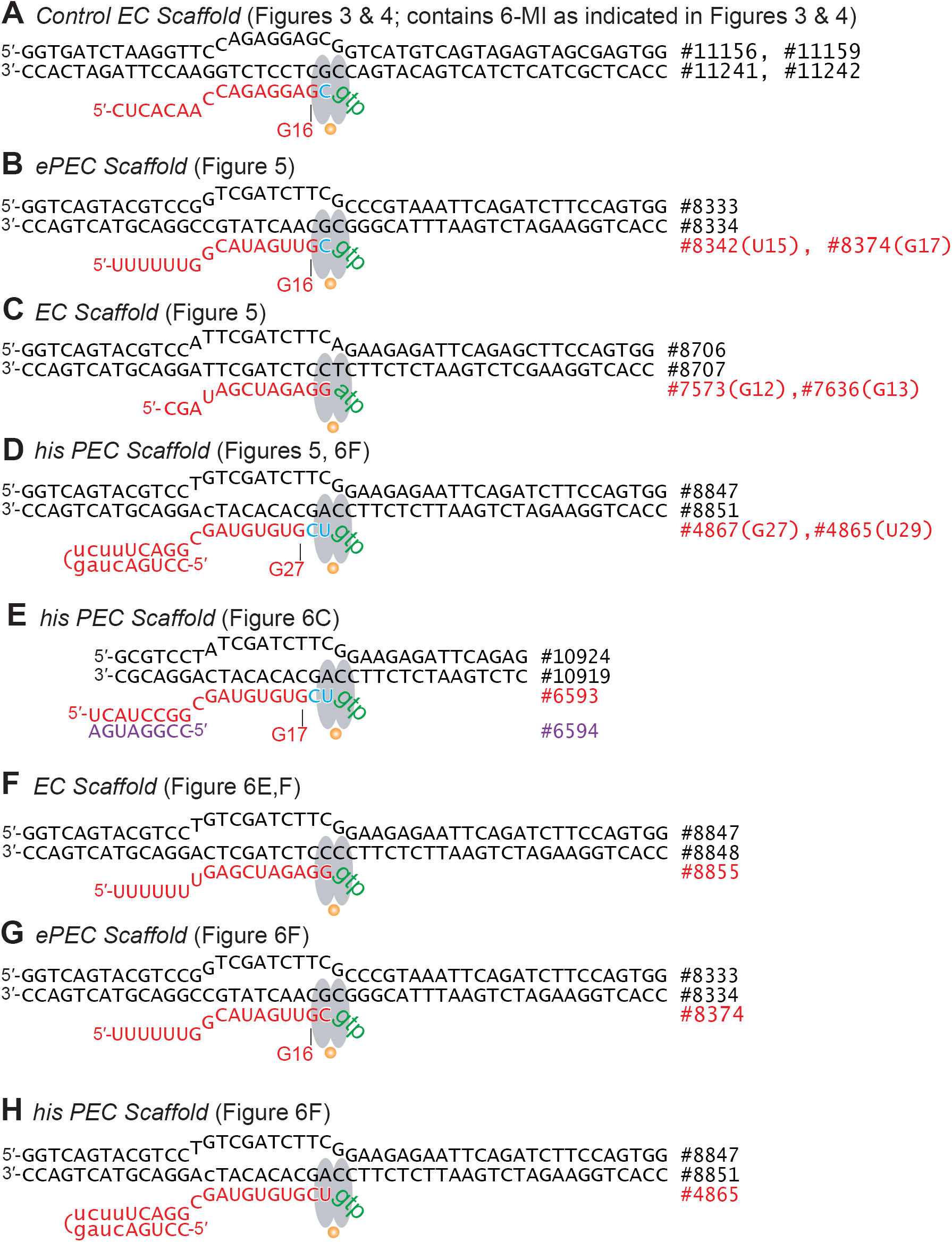
Scaffolds used for experiments, if not shown in other figures. (**A**) The control EC scaffold for 6-MI translocation assays in Figures 3 and 4. 6-MI is present at the locations indicated in Figures 3 and 4. Full sequences of oligonucleotides are given in Table S1. (**B-D**) Scaffolds used for disulfide bond assays of trigger loop position shown in Figure 5. For experiments on RNA 3’ OH-containing EC and PECs (Figures 5B-D), the full-length RNAs were used. For experiments on 3’deoxyEC and 3’deoxyPECs (Figures 5E,F), RNAs lacking the 3’ nucleotide were used for reconstitution and then extended with a 3’dNMP by RNAP-mediated catalysis. (**E**) *his*PEC scaffold used in Figure 5C (ntDNA#10924, tDNA #10919, RNA #6593+#6594; Table S1). In this scaffold, the *his* pause hairpin is mimicked by a 8-bp duplex that stabilizes the pause indistinguishably from the natural *his* pause hairpin (Hein *et al*., 2014). (**F**) Control EC scaffold used in Figure 6E,F (ntDNA#8847, tDNA #8848, RNA #8855; Table S1). (**G**) ePEC scaffold used in Figure 6F (ntDNA#8833, tDNA #8834, RNA #8374; Table S1). (**H**) *his*PEC scaffold used in Figure 6F (ntDNA#8847, tDNA #8851, RNA #4865; Table S1).

## REFERENCES

Aivazashvili, V. A., Bibilashvili, R., Vartikian, R. M. & Kutateladze, T. A. 1981. [Effect of the primary structure of RNA on the pulse character of RNA elongation in vitro by Escherichia coli RNA polymerase: a model]. Mol Biol (Mosk), 15, 915–29.

Artsimovitch, I. & Landick, R. 2002. The transcriptional regulator RfaH stimulates RNA chain synthesis after recruitment to elongation complexes by the exposed nontemplate DNA strand. Cell, 109, 193–203.

Bai, L., Shundrovsky, A. & Wang, M. D. 2004. Sequence-dependent kinetic model for transcription elongation by RNA polymerase. J Mol Biol, 344, 335–49.

Bochkareva, A., Yuzenkova, Y., Tadigotla, V. R. & Zenkin, N. 2012. Factor-independent transcription pausing caused by recognition of the RNA-DNA hybrid sequence. EMBO J, 31, 630–9.

Brueckner, F. & Cramer, P. 2008. Structural basis of transcription inhibition by alpha-amanitin and implications for RNA polymerase II translocation. Nat Struct Mol Biol, 15, 811–8.

Chan, C., Wang, D. & Landick, R. 1997. Spacing from the transcript 3’ end determines whether a nascent RNA hairpin interacts with RNA polymerase to prolong pausing or triggers termination. J. Mol. Biol., 268, 54–68.

Chan, C. L. & Landick, R. 1993. Dissection of the *his* leader pause site by base substitution reveals a multipartite signal that includes a pause RNA hairpin. J. Mol. Biol., 233, 25–42.

Chen, H., Shiroguchi, K., Ge, H. & Xie, X. S. 2015. Genome-wide study of mRNA degradation and transcript elongation in Escherichia coli. Mol Syst Biol, 11, 781.

Churchman, L. S. & Weissman, J. S. 2012. Native elongating transcript sequencing (NET-seq). Curr Protoc Mol Biol, Chapter 4, Unit 4 14 1-17.

Dangkulwanich, M., Ishibashi, T., Liu, S., Kireeva, M. L., Lubkowska, L., Kashlev, M. & Bustamante, C. J. 2013. Complete dissection of transcription elongation reveals slow translocation of RNA polymerase II in a linear ratchet mechanism. Elife, 2, e00971.

Forde, N. R., Izhaky, D., Woodcock, G. R., Wuite, G. J. & Bustamante, C. 2002. Using mechanical force to probe the mechanism of pausing and arrest during continuous elongation by Escherichia coli RNA polymerase. Proc Natl Acad Sci U S A, 99, 11682–7.

Frank, J. 2017. Time-resolved cryo-electron microscopy: Recent progress. J Struct Biol, 200, 303–306.

Gabizon, R., Lee, A., Vahedian-Movahed, H., Ebright, R. H. & Bustamante, C. J. 2018. Pause sequences facilitate entry into long-lived paused states by reducing RNA polymerase transcription rates. Nat Commun, 9, 2930–2940.

Galburt, E. A., Grill, S. W., Wiedmann, A., Lubkowska, L., Choy, J., Nogales, E., Kashlev, M. & Bustamante, C. 2007. Backtracking determines the force sensitivity of RNAP II in a factor-dependent manner. Nature, 446, 820–3.

Gilbert, W., Maizels, N. & Maxam, A. 1974. Sequences of controlling regions of the lactose operon. Cold Spring Harbor Symp. Quant. Biol., 38, 845–855.

Guo, X., Myasnikov, A. G., Chen, J., Crucifix, C., Papai, G., Takacs, M., Schultz, P. & Weixlbaumer, A. 2018. Structural basis for NusA stabilized transcriptional pausing. Mol Cell, 69, 816-827 e4.

Gusarov, I. & Nudler, E. 1999. The mechanism of intrinsic transcription termination. Mol. Cell, 3, 495–504.

Hein, P. P., Kolb, K. E., Windgassen, T., Bellecourt, M. J., Darst, S. A., Mooney, R. A. & Landick, R. 2014. RNA polymerase pausing and nascent-RNA structure formation are linked through clamp-domain movement. Nat Struct Mol Biol, 21, 794–802.

Herbert, R., La Porta, A., Wong, B., Mooney, R., Neuman, K., Landick, R. & Block, S. 2006. Sequence-resolved detection of pausing by single RNA polymerase molecules. Cell, 125, 1083–1094.

Hubin, E. A., Fay, A., Xu, C., Bean, J. M., Saecker, R. M., Glickman, M. S., Darst, S. A. & Campbell, E. A. 2017. Structure and function of the mycobacterial transcription initiation complex with the essential regulator RbpA. Elife, 6, e22520.

Imashimizu, M., Kireeva, M. L., Lubkowska, L., Gotte, D., Parks, A. R., Strathern, J. N. & Kashlev, M. 2013. Intrinsic translocation barrier as an initial step in pausing by RNA polymerase II. J Mol Biol, 425, 697–712.

Imashimizu, M., Takahashi, H., Oshima, T., Mcintosh, C., Bubunenko, M., Court, D. L. & Kashlev, M. 2015. Visualizing translocation dynamics and nascent transcript errors in paused RNA polymerases in vivo. Genome Biol, 16, 98.

Johnson, K. A., Simpson, Z. B. & Blom, T. 2009. Global kinetic explorer: a new computer program for dynamic simulation and fitting of kinetic data. Anal Biochem, 387, 20–9.

Jonkers, I. & Lis, J. T. 2015. Getting up to speed with transcription elongation by RNA polymerase II. Nat Rev Mol Cell Biol, 16, 167–77.

Kamarthapu, V., Epshtein, V., Benjamin, B., Proshkin, S., Mironov, A., Cashel, M. & Nudler, E. 2016. ppGpp couples transcription to DNA repair in E. coli. Science, 352, 993–6.

Kang, J. Y., Mishanina, T. V., Bellecourt, M. J., Mooney, R. A., Darst, S. A. & Landick, R. 2018a. RNA polymerase accommodates a pause RNA hairpin by global conformational rearrangements that prolong pausing. Mol Cell, 69, 802–815 e1.

Kang, J. Y., Mooney, R. A., Nedialkov, Y., Saba, J., Mishanina, T. V., Artsimovitch, I., Landick, R. & Darst, S. 2018b. Structural basis for transcript elongation control by NusG/RfaH universal regulators. Cell, 137, 1650–1662.

Kang, J. Y., Olinares, P. D., Chen, J., Campbell, E. A., Mustaev, A., Chait, B. T., Gottesman, M. E. & Darst, S. A. 2017. Structural basis of transcription arrest by coliphage HK022 nun in an Escherichia coli RNA polymerase elongation complex. Elife, 6, e25478.

Kassavetis, G. A. & Chamberlin, M. J. 1981. Pausing and termination of transcription within the early region of bacteriophage T7 DNA in vitro. J Biol Chem, 256, 2777–86.

Kingston, R. E., Nierman, W. C. & Chamberlin, M. J. 1981. A direct effect of guanosine tetraphosphate on pausing of Escherichia coli RNA polymerase during RNA chain elongation. J Biol Chem, 256, 2787–97.

Kireeva, M. L. & Kashlev, M. 2009. Mechanism of sequence-specific pausing of bacterial RNA polymerase. Proc Natl Acad Sci U S A, 106, 8900–5.

Kyzer, S., Ha, K. S., Landick, R. & Palangat, M. 2007. Direct versus limited-step reconstitution reveals key features of an RNA hairpin-stabilized paused transcription complex. J Biol Chem, 282, 19020–8.

Landick, R. 2006. The regulatory roles and mechanism of transcriptional pausing. Biochem Soc Trans, 34, 1062–6.

Landick, R., Carey, J. & Yanofsky, C. 1985. Translation activates the paused transcription complex and restores transcription of the *trp* operon leader region. Proc. Natl. Acad. Sci. USA, 82, 4663–4667.

Larson, M. H., Mooney, R. A., Peters, J. M., Windgassen, T., Nayak, D., Gross, C. A., Block, S. M., Greenleaf, W. J., Landick, R. & Weissman, J. S. 2014. A pause sequence enriched at translation start sites drives transcription dynamics in vivo. Science, 344, 1042–7.

Liu, B., Zuo, Y. & Steitz, T. A. 2016. Structures of E. coli sigmaS-transcription initiation complexes provide new insights into polymerase mechanism. Proc Natl Acad Sci U S A, 113, 4051–6.

Malinen, A. M., Nandymazumdar, M., Turtola, M., Malmi, H., Grocholski, T., Artsimovitch, I. & Belogurov, G. A. 2014. CBR antimicrobials alter coupling between the bridge helix and the beta subunit in RNA polymerase. Nat Commun, 5, 3408.

Malinen, A. M., Turtola, M., Parthiban, M., Vainonen, L., Johnson, M. S. & Belogurov, G. A. 2012. Active site opening and closure control translocation of multisubunit RNA polymerase. Nucleic Acids Res, 40, 7442–51.

Maoileidigh, D. O., Tadigotla, V. R., Nudler, E. & Ruckenstein, A. E. 2011. A unified model of transcription elongation: what have we learned from single-molecule experiments? Biophys J, 100, 1157–66.

Mayer, A., Landry, H. M. & Churchman, L. S. 2017. Pause & go: from the discovery of RNA polymerase pausing to its functional implications. Curr Opin Cell Biol, 46, 72–80.

Mejia, Y. X., Nudler, E. & Bustamante, C. 2014. Trigger loop folding determines transcription rate of Escherichia coli’s RNA polymerase. Proc Natl Acad Sci U S A.

Mishanina, T. V., Palo, M. Z., Nayak, D., Mooney, R. A. & Landick, R. 2017. Trigger loop of RNA polymerase is a positional, not acid-base, catalyst for both transcription and proofreading. Proc Natl Acad Sci U S A, 114, E5103–E5112.

Murakami, K. S., Shin, Y., Turnbough, C. L., Jr. & Molodtsov, V. 2017. X-ray crystal structure of a reiterative transcription complex reveals an atypical RNA extension pathway. Proc Natl Acad Sci U S A.

Nayak, D., Voss, M., Windgassen, T., Mooney, R. A. & Landick, R. 2013. Cys-pair reporters detect a constrained trigger loop in a paused RNA polymerase. Mol Cell, 50, 882–93.

Palangat, M., Hittinger, C. T. & Landick, R. 2004. Downstream DNA selectively affects a paused conformation of human RNA polymerase II. J. Mol. Biol., 341, 429–442.

Palangat, M. & Landick, R. 2001. Roles of RNA:DNA hybrid stability, RNA structure, and active site conformation in pausing by human RNA polymerase II. J Mol Biol, 311, 265–82.

Pan, T., Artsimovitch, I., Fang, X. W., Landick, R. & Sosnick, T. R. 1999. Folding of a large ribozyme during transcription and the effect of the elongation factor NusA. Proc Natl Acad Sci U S A, 96, 9545–50.

Pan, T. & Sosnick, T. 2006. RNA folding during transcription. Annu Rev Biophys Biomol Struct, 35, 161–75.

Proshkin, S., Rahmouni, A. R., Mironov, A. & Nudler, E. 2010. Cooperation between translating ribosomes and RNA polymerase in transcription elongation. Science, 328, 504–8.

Proudfoot, N. J. 2016. Transcriptional termination in mammals: Stopping the RNA polymerase II juggernaut. Science, 352, aad9926.

Sekine, S., Murayama, Y., Svetlov, V., Nudler, E. & Yokoyama, S. 2015. The ratcheted and ratchetable structural states of RNA polymerase underlie multiple transcriptional functions. Mol Cell, 57, 408–21.

Shu, B. & Gong, P. 2016. Structural basis of viral RNA-dependent RNA polymerase catalysis and translocation. Proc Natl Acad Sci U S A, 113, E4005–14.

Siebert, M. & Söding, J. 2016. Bayesian Markov models consistently outperform PWMs at predicting motifs in nucleotide sequences. Nucleic Acids Res, 44, 6055–69.

Sosunova, E., Sosunov, V., Epshtein, V., Nikiforov, V. & Mustaev, A. 2013. Control of transcriptional fidelity by active center tuning as derived from RNA polymerase endonuclease reaction. J Biol Chem, 288, 6688–703.

Steinert, H., Sochor, F., Wacker, A., Buck, J., Helmling, C., Hiller, F., Keyhani, S., Noeske, J., Grimm, S., Rudolph, M. M., Keller, H., Mooney, R. A., Landick, R., Suess, B., Furtig, B., Wohnert, J. & Schwalbe, H. 2017. Pausing guides RNA folding to populate transiently stable RNA structures for riboswitch-based transcription regulation. Elife, 6.

Strobel, E. J. & Roberts, J. W. 2015. Two transcription pause elements underlie a sigma70-dependent pause cycle. Proc Natl Acad Sci U S A, 112, E4374–80.

Tadigotla, V. R., D, O. M., Sengupta, A. M., Epshtein, V., Ebright, R. H., Nudler, E. & Ruckenstein, A. E. 2006. Thermodynamic and kinetic modeling of transcriptional pausing. Proc Natl Acad Sci U S A, 103, 4439–44.

Toulokhonov, I., Artsimovitch, I. & Landick, R. 2001. Allosteric control of RNA polymerase by a site that contacts nascent RNA hairpins. Science, 292, 730–3.

Toulokhonov, I., Zhang, J., Palangat, M. & Landick, R. 2007. A central role of the RNA polymerase trigger loop in active-site rearrangement during transcriptional pausing. Mol Cell, 27, 406–19.

Vassylyev, D., Vassylyeva, M., Perederina, A., Tahirov, T. & Artsimovitch, I. 2007. Structural basis for transcription elongation by bacterial RNA polymerase. Nature, 448, 157–162.

Vvedenskaya, I. O., Vahedian-Movahed, H., Bird, J. G., Knoblauch, J. G., Goldman, S. R., Zhang, Y., Ebright, R. H. & Nickels, B. E. 2014. Transcription. Interactions between RNA polymerase and the “core recognition element” counteract pausing. Science, 344, 1285–9.

Wang, D., Bushnell, D. A., Huang, X., Westover, K. D., Levitt, M. & Kornberg, R. D. 2009. Structural basis of transcription: backtracked RNA polymerase II at 3.4 angstrom resolution. Science, 324, 1203–6.

Wang, D., Meier, T., Chan, C., Feng, G., Lee, D. & Landick, R. 1995. Discontinuous movements of DNA and RNA in E. coli RNA polymerase accompany formation of a paused transcription complex. Cell, 81, 341–350.

Weixlbaumer, A., Leon, K., Landick, R. & Darst, S. A. 2013. Structural basis of transcriptional pausing in bacteria. Cell, 152, 431–41.

Wickiser, J. K., Winkler, W. C., Breaker, R. R. & Crothers, D. M. 2005. The speed of RNA transcription and metabolite binding kinetics operate an FMN riboswitch. Mol Cell, 18, 49–60.

Windgassen, T., Mooney, R. A., Nayak, D., Palangat, M., Zhang, J. & Landick, R. 2014. Trigger-helix folding pathway and SI3 mediate catalysis and hairpin-stabilized pausing by *Escherichia coli* RNA polymerase. Nucleic Acids Res, 42, 12707–21.

Yakhnin, A. V., Murakami, K. S. & Babitzke, P. 2016. NusG Is a Sequence-specific RNA Polymerase Pause Factor That Binds to the Non-template DNA within the Paused Transcription Bubble. J Biol Chem, 291, 5299–308.

Zhang, J. & Landick, R. 2016. A two-way street: Regulatory interplay between RNA polymerase and nascent RNA structure. Trends Biochem Sci, 41, 293–310.

Zhang, J., Palangat, M. & Landick, R. 2010. Role of the RNA polymerase trigger loop in catalysis and pausing. Nat Struct Mol Biol, 17, 99–104.

